# Single-cell RNA sequencing reveals distinct senotypes and a quiescence-senescence continuum at the transcriptome level following chemotherapy

**DOI:** 10.1101/2025.05.13.653730

**Authors:** Brianna Fernandez, Victor Passanisi, Humza M. Ashraf, Sabrina L. Spencer

## Abstract

Quiescence (reversible cell-cycle arrest) and senescence (irreversible arrest) are challenging to distinguish due to a lack of specific biomarkers, yet both arise simultaneously after chemotherapy. While senescence suppresses tumors by limiting proliferation and recruiting the immune system, quiescent cancer cells evade future therapies and may resume proliferation. Here, we pair time-lapse imaging of cell-cycle dynamics with single-cell RNA-sequencing after etoposide treatment to differentiate these states, linking heterogeneous cell-cycle phenotypes to the transcriptomic landscape. We identify diverse senescent types (senotypes) and link them to two arrest pathways – a gradual path arising after a standard mitosis-to-G0 transition, and a rarer but direct path driven by a mitotic slip. Using pseudotime trajectory analysis, we find that senescent phenotypes begin to manifest early and gradually along the first trajectory, even in shallow quiescent cells. These data support a model wherein, following chemotherapy, quiescence and senescence exist on a continuum of cell-cycle withdrawal at a transcriptome-wide level.

## Introduction

The cellular decision to proliferate or withdraw from the cell cycle is essential for maintaining tissue health and homeostasis. When this decision goes awry, it results in diseases of hypo-proliferation, such as aging, or hyper-proliferation, such as cancer. Despite its importance, the molecular details of cell-cycle withdrawal are not fully understood. Quiescence is a state of transient and reversible cell-cycle withdrawal that occurs in response to low levels of DNA damage or lack of mitogens or nutrients^1^. Senescence is a state of irreversible cell-cycle withdrawal wherein cells remain metabolically active and viable but never divide again^2,3^. Senescence has a normal, physiological role in development, but also occurs in response to sublethal DNA-damage or stress^4^. This arrest is initially triggered and maintained by the p53/p21 and p16/Rb signaling axes^2,3,5^ and is accompanied by many cellular changes including increases in cell size, autophagy, changes in chromatin, and the secretion of pro-inflammatory and pro-proliferation proteins called the senescence associated secretory phenotype (SASP)^2,6–8^.

Therapy induced senescence (TIS) is a permanent cell-cycle withdrawal induced by exposure to radiation or chemotherapeutic drugs like etoposide and doxorubicin^9,10^. In the context of cancer, TIS has been identified as an alternative therapeutic strategy to cell death since these cells are irreversibly arrested, preventing the proliferation of damaged premalignant cells and initiating immune clearance through the SASP^9,11^. There is still debate about TIS as an end goal for therapies because some work suggests that some cells may be able to evade TIS, remaining dormant for a time before returning to the cell cycle^12–14^. Therefore, understanding the molecular relationship between cells that are transiently quiescent and cells that are senescent is critically important.

Studying the relationship between these different arrested states is practically challenging due to the lack of distinct, consistent, and specific biomarkers^15^, and because measuring irreversible arrest in heterogeneous populations requires single-cell, time-lapse data to define the ground-truth senescent cells. To overcome this, we recently used single-cell time-lapse microscopy and a live-cell CDK2 activity reporter to track each cell’s cumulative cell-cycle withdrawal duration across thousands of single cells recovering from etoposide, oxidative stress, or ionizing radiation^16^. This approach distinguished irreversibly arrested senescent cells from slow-cycling cells that spend long periods in quiescence between cell cycles and showed that both fates arise from the same treatment. By correlating each cell’s withdrawal duration with the intensity of eight canonical senescence markers, we found that both quiescent and senescent cells express these eight markers in a graded fashion. Furthermore, the graded intensity of each marker across the population reflected each cell’s duration of cell-cycle withdrawal^16^. These findings suggested that chemotherapy-induced cell-cycle withdrawal may be a graded continuum rather than a binary decision between quiescence and senescence. However, an open question following this work is whether binary markers of quiescence vs. senescence might nevertheless exist, and whether these eight graded markers are representative of changes across the entire transcriptome. Put another way, at a transcriptome level, it is unclear whether quiescent and senescent states exist as distinct islands within a population of heterogeneous cells, or whether they would exist on a continuum and be connected by a bridge.

Here, we addressed these questions by measuring the changing transcriptome in senescent cells by performing live-cell time-lapse microscopy in parallel with single-cell RNA sequencing in MCF10A mammary epithelial cells following chemotherapy. This approach avoids reliance on senescence markers because it uses long-term cell-cycle withdrawal to define senescent cells, and enabled transcriptome-wide analysis of gene expression changes. We uncovered a gradient of cellular states corresponding to varying probabilities of cell-cycle re-entry, supporting the quiescence-senescence continuum model at a transcriptome-wide level. Notably, quiescent cells lacked a unique signature and instead exhibited attenuated senescence-associated gene expression changes. Additionally, within a single population, we identified distinct senescence types, or “senotypes”, with distinct gene expression profiles that resulted from different paths to senescence. This work contributes to a growing body of research that aims to fully understand the molecular characteristics associated with reversible and irreversible cell-cycle withdrawal.

## Results

### Establishment of populations with increasing fractions of senescent cells

To study senescent cells at the transcriptomic level, it is essential to identify experimental conditions that yield a homogeneous senescent population as measured by a “ground-truth” assessment of senescence: the total lack of proliferation over time. To test whether we could alter the proportion of quiescent and senescent cells, we treated MCF10A cells with increasing doses of the chemotherapy etoposide, a widely used inducer of senescence that causes DNA damage and cell-cycle arrest^17^. After treating cells for 24 h, we washed off the etoposide and allowed cells to recover in drug-free media for 6 days (Fig. 1A). On day 6, cells were fixed and stained for phospho-Rb, a marker of cell-cycle commitment^18^ (Fig. 1B). Higher etoposide doses resulted in fewer cycling (pRb^high^) cells. In parallel, we used time-lapse microscopy to track MCF10A cells expressing a DHB-based CDK2 activity reporter^19^ (Fig. S1A) and an H2B-mTq nuclear marker from day 6 to day 10 post etoposide. CDK2 activity turns on at cell-cycle commitment, rising throughout the cell cycle, and turns off when cells withdraw from the cell cycle. Higher etoposide doses produced a greater proportion of CDK2^low^ non-cycling cells over the 4-day imaging period (Fig. 1C-D). We plotted the distribution of cell-cycle withdrawal durations, measured as time spent CDK2^low^ (CDK2 activity < 0.8) for each etoposide dose and categorized cells as fast-cycling, slow-cycling, or predicted-senescent based on CDK2^low^ durations (Fig. 1E and Fig. S1B-C). At 2.5µM and 10µM, we observed a mix of all three categories, while 25µM resulted in predominantly senescent cells. We confirmed the expected expression of canonical senescent markers using immunofluorescent (IF) imaging immediately following the live-cell movies (Fig. S1D). These results show that etoposide dose modulates the proportion of predicted-senescent cells as well as their expression of canonical senescent markers.

**Figure 1.**
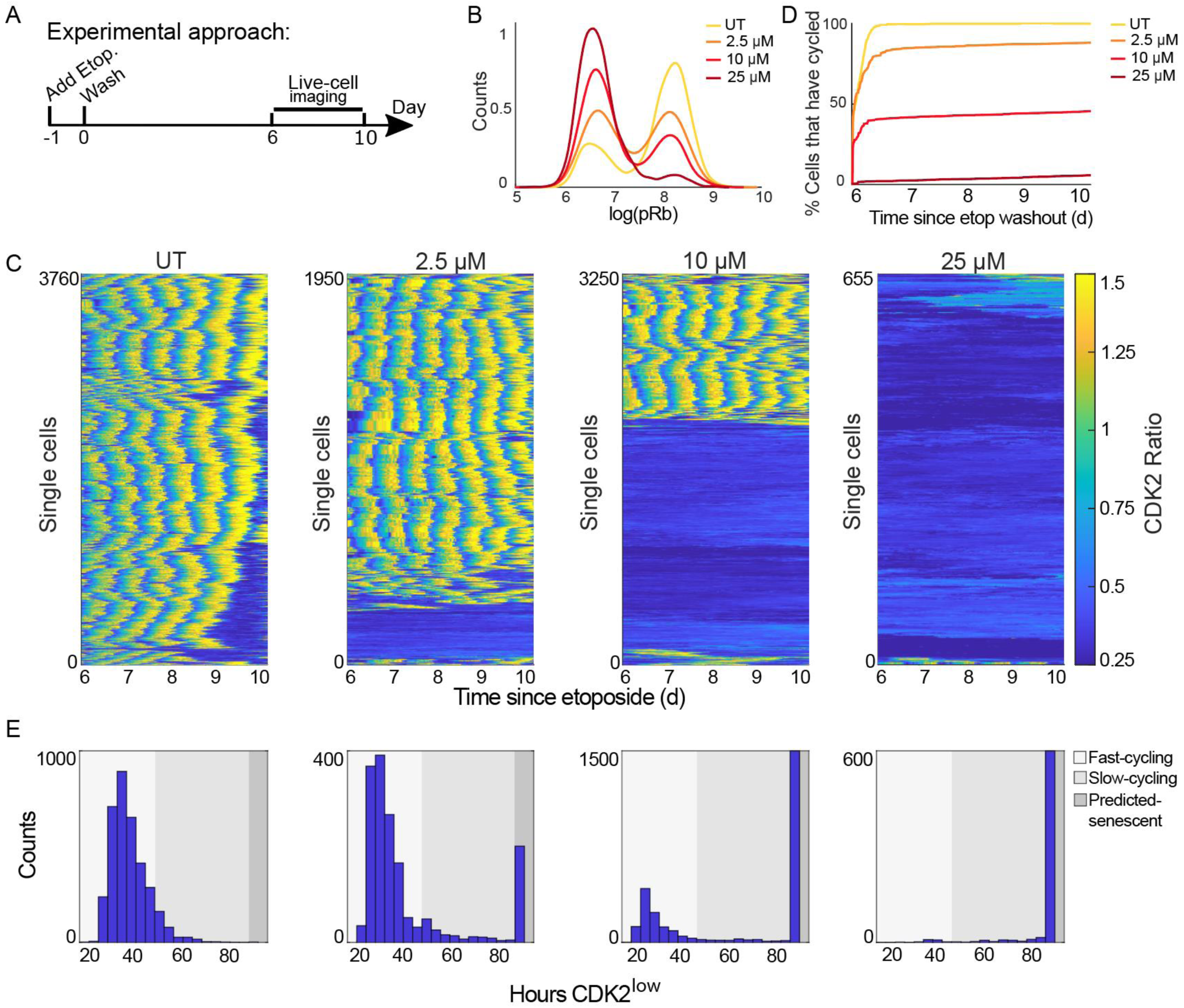
Increasing the dose of etoposide increases the proportion of predicted-senescent cells. **(A)** Experimental timeline. **(B)** MCF10A cells were treated with 2.5 μM, 10μM, or 25μM etoposide for 24h followed by a 6d drug-free recovery, at which point cells were stained for phospho-Rb (S807/811) to measure the fraction of cycling (pRb^high^) and non-cycling (pRb^low^) cells, visualized as a histogram. Log refers to natural log and UT refers to untreated throughout this work. **(C)** MCF10A cells expressing the CDK2 activity sensor were treated as in A. Cells were filmed by fluorescent time-lapse microscopy for 96h, from 6d-10d post etoposide. Each row in the heatmap represents a single-cell trace, colored according to the colormap where yellow indicates high CDK2 activity and progression through the cell cycle, and dark blue indicates cell-cycle withdrawal into quiescence or senescence. **(D)** Percent of cells that have entered the cell cycle (CDK2 activity > 0.8) at each frame of the movie in C. **(E)** Distributions of CDK2^low^ times (CDK2 activity < 0.8) for the movie in C. Shading represents the CDK2^low^ times corresponding to fast-cycling, slow-cycling, and non-cycling categories. Throughout the paper, the number of cells plotted and number of biological replicates can be found in Supplementary table 2.

### Transcriptomic profiling of etoposide-treated cells reveals two paths to senescence and diverse senescent types

To understand the transcriptional changes that occur as cells move between the different cell-cycle fates we observed by live-cell imaging, we performed single-cell RNA sequencing of cells at each of the etoposide doses. We sequenced MCF10A cells 6d after release from a 24h treatment with 2.5µM, 10µM, or 25µM etoposide (Fig 2A and S2A) using the 10x Genomics protocol. Cells with low read counts and genes were filtered out (S2B, see methods) resulting in 6454, 5913, 5361, and 4878 cells profiled for UT, 2.5µM, 10µM, and 25µM etoposide, respectively.

**Figure 2.**
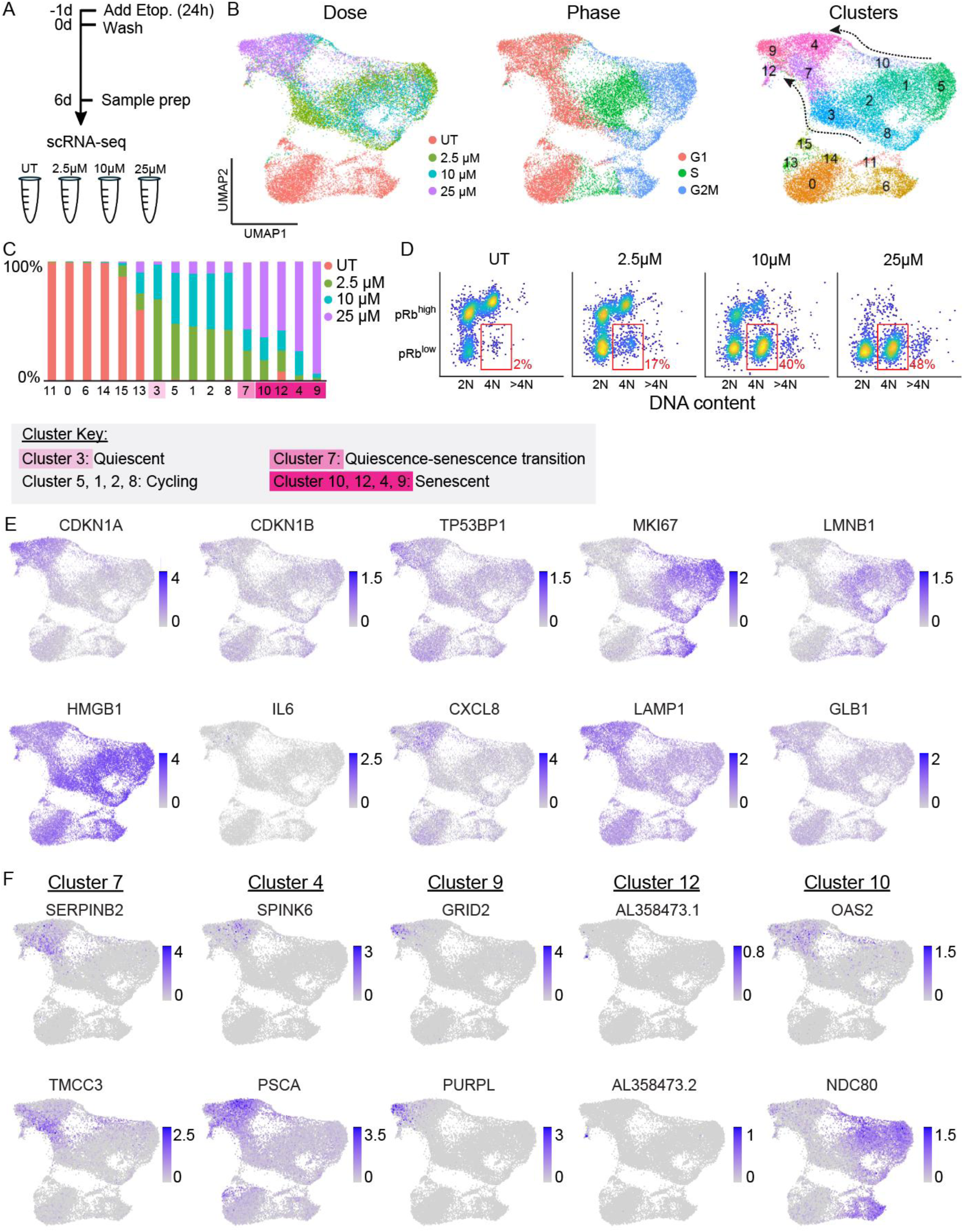
Transcriptomic profiling of etoposide-treated cells. **(A)** Experimental timeline. **(B)** UMAP projection of scRNA-seq results with cells colored by etoposide dose (left), cell-cycle phase according to Seurat (middle), and clustered by gene expression similarity, where arrows depict two paths or bridges to senescence (right). **(C)** Proportion of each cluster in B by etoposide dose, and key with interpretation of the cell-cycle state of each cluster. **(D)** MCF10A cells were treated with 2.5 μM, 10μM, or 25μM etoposide for 24h followed by a 6d drug-free recovery. Cells were stained for phospho-Rb (S807/811) and Hoechst for DNA content and plotted as a density scatter where yellow indicates a large number of cells in that region. The same number of cells is plotted for each dose. Red box indicates the pRb^low^ /4N cells, a hallmark of cells that have experienced a mitotic slip. **(E)** UMAP projection colored by normalized and scaled expression (see methods) for 10 canonically used senescent markers. **(F)** UMAP projection colored by normalized and scaled expression for two top single-gene markers (ranked by log2FC * (-log(p.adjusted))) from cluster 7 (quiescence-senescence transition cluster) and clusters 4, 9, 12, and 10 (senescent clusters).

We used Uniform Manifold Approximation and Projection (UMAP)^20^ to project all cells together in the same 2-dimensional space (Fig 2B). Untreated cells formed the lower island of the UMAP, while the three different doses of etoposide-treated cells formed the upper island. Cells treated with 25µM etoposide occupied a region on the far-left side of the upper island, whereas 2.5µM and 10µM cells were dispersed along the upper island (Fig 2B left and S2C).

We used Seurat^21,22^ to categorize cells as either G1, S, or G2/M based on the expression of cell-cycle phase-specific genes (Fig 2B middle and S2D, see methods). Seurat does not distinguish between cycling G1 phase cells and G0 phase quiescent cells – all cells that are not expressing S or G2/M markers are grouped together and labeled “G0/G1” and will contain both quiescent and senescent cells. As expected based on our live-cell imaging, 25µM etoposide cells that we know are predominantly senescent were labelled as G0/G1, whereas 2.5µM and 10µM cells were labelled as a mix of G0/G1, S, and G2/M (Fig 2B middle).

Notably, untreated S and G2/M cells were transcriptionally distinct from S and G2/M etoposide-treated cells and occupied different regions of the UMAP (Fig 2B middle), despite having similar CDK2 activities when measured by live-cell imaging (Fig 1C). This suggests that while some cells can evade the proliferative arrest from lower etoposide doses and continue to cycle, they retain an altered transcriptional state even 6d after the drug is washed off. Cells that cycle after etoposide treatment showed increased expression of protein chaperones (Fig. S2H) when compared to untreated cycling cells, a possible adaptive mechanism allowing them to cycle with residual etoposide-induced stress, and a potential weakness to exploit to target cycling drug-tolerant persister cells in cancer.

We clustered cells by similarity (see methods) and measured the number of cells in each cluster from each dose of etoposide (Fig. 2B right and Fig. 2C). Most clusters in the treated island of the UMAP were composed of cells from more than one dose of etoposide, reflecting the heterogeneity in cell-cycle fate we saw in our live-cell imaging for 2.5µM and 10µM etoposide. By contrast, 25µM cells fell into 5 different clusters on the left side of the UMAP, suggesting transcriptional heterogeneity within the predicted-senescent cells themselves.

Based on the cell-cycle behavior measured by live-cell imaging, we identified clusters 9, 12, 4, and 10 as predicted-senescent clusters because they 1) they consisted primarily of 25µM cells that we know from time-lapse imaging to be senescent, and 2) showed increasing cluster occupancy as a function of etoposide dose (S2C). The nearby cluster 7 contained a mix of 2.5µM, 10µM, and 25µM etoposide cells, suggesting that it represents a quiescent-senescent transition cluster. Cluster 3 is also comprised of cells with a G0/G1 label but has almost no 25µM cells and connects to the cycling cells, likely representing a cluster of quiescent cells.

Importantly, we found two different “bridges” between the etoposide-treated S/G2/M cycling cells and the senescent region of the UMAP (Fig 2B). Clusters 3 and 7 made up the dominant bridge and included primarily 2.5µM and 10µM etoposide cells; cluster 10 formed a less populated alternative bridge and primarily consisted of 25µM cells (arrows in Fig 2B right). We hypothesized that these two paths represented, respectively, a standard G0/G1 arrest after completion of mitosis, and a mitotic slip following G2 arrest. A mitotic slip causes a cell to enter a G0/G1-like state without mitosis and yields a population of cells with 4N DNA content^23^ and elevated levels of the CDK inhibitor, p21^24^. To validate that the cluster 10 “bridge” indeed represented cells that had undergone a mitotic slip, we co-stained cells for pRb and DNA content at each dose. We detected 2N DNA content/pRb^low^ cells (standard completion of mitosis) and an increasing proportion of 4N DNA content/pRb^low^ (mitotic slippage) cells with increasing etoposide dose (Fig 2D). Identification of cells that have experienced a mitotic slip via scRNA-seq is notable because this represents a direct path from cycling (cluster 5) to senescence (cluster 10), without transiting through quiescence.

We next visualized the expression of ten canonical senescence biomarkers across all cells (Fig 2E and S2F). While specific marker expression patterns varied, the canonical markers of senescence fail to specifically identify senescent cells at the mRNA level, with the exception of CDKN1A (p21), which showed a marked, albeit graded, increase in all senescent clusters.

While multiple studies of senescence exist, the extent to which individual cells within the same population are transcriptionally different is less well studied. To determine the specific differences in expression profiles of our senescent clusters, we performed differential gene expression (DGE) analysis on each senescent cluster compared to all other cells. DGE analysis revealed specifically upregulated markers for cluster 7, cluster 4, cluster 9, and cluster 12 that could be used to light up these clusters reasonably uniquely (Fig 2F and S2F right). Cluster 9 was particularly well-marked by upregulation of PURPL, a p53-induced lncRNA with emerging roles in senescence and cancer^25–28^, and by GRID2, an ionotropic receptor involved in glutamate transfer. Cluster 12 was particularly well marked by increased expression of many lncRNAs. However, cluster 10 genes showed less cluster-specific upregulation. This analysis thus revealed several genes whose distinctive expression can be used to flag the senescent clusters we identified. By contrast, the top downregulated genes were much less specific to each cluster (Fig S2E and S2F left).

To get a high-level view of expression patterns in our senescent cells, we plotted the expression of the top 50 up- and down-regulated genes for each of the senescent clusters in all pathways (Fig S2G). Generally, up-regulated genes showed more cluster specificity whereas down-regulated genes were either down in all etoposide treated cells regardless of cell-cycle status, down in all non-cycling cells including untreated cells, or down in all senescent clusters.

### Pathway analysis reveals biological processes altered in senescent cells

To determine if the most differentially expressed (DE) genes for each senescent cluster (Fig S3A) belonged to a shared biological process, we performed gene set enrichment analysis (GSEA)^29^ using the Gene Ontology (GO) Biological Processes gene set annotations^30^ (see methods). We found multiple significantly upregulated (p.adjusted < 0.05) and downregulated pathways for clusters 7, 4, 9, and 10 (Fig 3A). Cluster 12 had no significant up- or downregulated pathways, since many of the cluster 12 DE genes were lncRNAs (S2F) and not well annotated in specific pathways.

**Figure 3.**
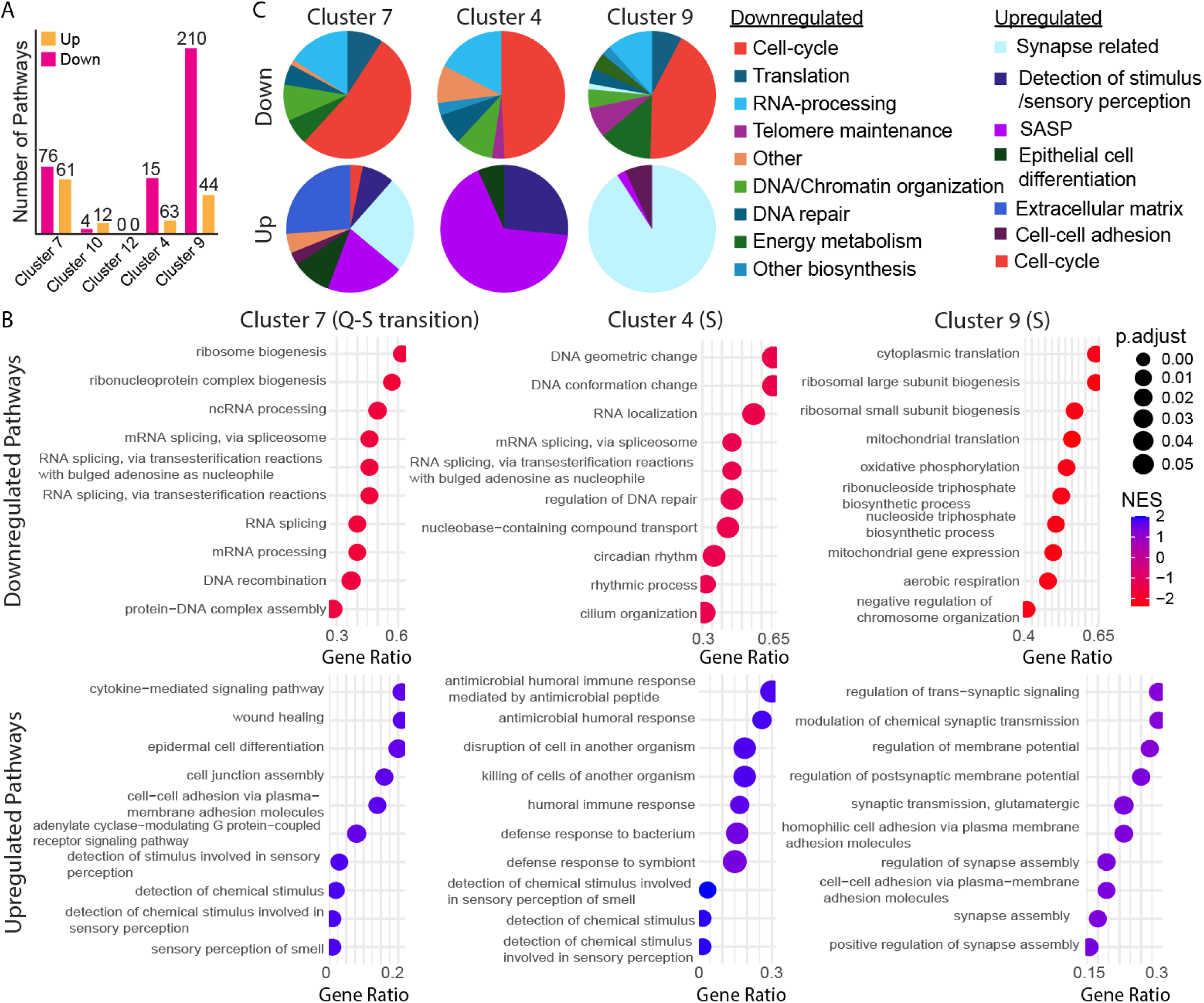
Pathway analysis reveals diverse types of senescent cells. **(A)** Number of pathways significantly up- and down-regulated for each cluster (p.adjusted < 0.05). **(B)** Significant pathways were grouped by general biological process (see methods, Supplementary table 1). **(C)** Gene set enrichment analysis (GSEA) for cluster 7 (quiescence-senescence transition cluster) and clusters 4 and 9 (senescent clusters). Top 10 pathways (by p.adjusted) for each cluster are displayed. Gene ratio for each pathway (proportion of genes in each pathway that are significantly DE compared to the total number of genes in that pathway) plotted on the x-axis. Dots colored by the normalized enrichment score for each pathway. Dot size reflects adjusted pvalue.

We plotted the gene ratio and normalized enrichment score (NES) for the top 10 (ranked by p.adjusted) up- and down-regulated pathways for each cluster, ignoring cell-cycle related pathways (Fig 3B and S3C). Many pathways had similar GO terms and overlapping leading-edge genes, so we grouped them based on general cellular process (see methods and Supplementary table 1) (Fig 3C). As expected, we found that pathways related to cell-cycle progression were the largest group of down-regulated pathways for all senescent clusters (Fig. 3C and S3B). Interestingly, the second and third largest groups of down-regulated pathways were related to the processing/transport of RNA and translation. We detected changes in these pathways in all senescent clusters.

Despite having DNA-damage from the etoposide treatment, cluster 9, 4, and 7 had decreased expression of genes associated with DNA repair pathways. This is likely because many DNA repair genes are E2F-target genes that are only expressed upon cell-cycle commitment. All three clusters also showed down-regulation of DNA/chromatin organization related pathways, indicating a shared feature of senescent cells. Cluster 4 had many downregulated genes belonging to the pathways for “DNA geometric change” and “DNA conformation change”, possibly due to the arrival in this state via mitotic slippage causing 4N DNA content. Cluster 9, the farthest away from cycling cells and the cluster with the highest p21 expression, had the most varied down-regulated pathway types, including translation, RNA processing, telomere maintenance, and energy/metabolism pathways.

We expected to see an increase in pathways associated with inflammation due to the SASP^7,8^, but we only saw this in clusters 4 and 7. Cluster 4 showed an increase in immune pathways such as “antimicrobial humoral response” and “defense response to bacterium”, whereas cluster 7 cells upregulated “wound healing” and “epidermal cell differentiation”. Notably, cluster 9 lacked SASP-related pathways and was instead defined by an upregulation of many synapse-related pathways including “regulation of trans-synaptic signaling” and “regulation of membrane potential”, with leading-edge genes that included many ion channels and metabolite sensing transmembrane receptors (Supplementary table 1).

Altogether, these data reveal that senescent cells exhibit gene expression changes associated with diverse biological processes. While some processes are shared, the extent to which they each manifest varies across clusters, leading to three senescent “types” within irreversibly arrested populations.

### scRNA-seq reveals a transcriptomic gradient along the quiescence-senescence continuum

If the continuum model of cell-cycle withdrawal is correct, then 1) quiescent cells would not have a unique gene expression program when compared to senescent cells and 2) quiescent cells would begin to show gene expression changes associated with late senescent phenotypes, even in G0 cells that are close to cycling.

To test if quiescent cells have a unique expression signature, we performed two DGE analyses. First, we compared quiescent cluster 3 to all other cells. The top up- and down-regulated DE genes for each cluster were not unique (Fig. S4A) and GSEA revealed that the most significantly down-regulated pathways were cell cycle-related (Fig. S4B). This suggests that the defining feature of quiescent cells when compared to all other cells is their lack of cycling markers. Second, we specifically compared quiescent and senescent cells by performing DGE analysis for quiescent cluster 3 vs. senescent clusters 4, 9, and 12, and transition cluster 7 vs. senescent clusters 4, 9, and 12. The most differentially expressed genes (Fig 4A) and pathways (Fig 4B) in cluster 3 vs. senescent cells were shared with cycling cells (cluster 2). These findings suggest that quiescent cells in the etoposide-released context studied here lack a unique transcriptional program and are instead defined by the lack of cycling genes and an attenuated senescent program.

**Figure 4.**
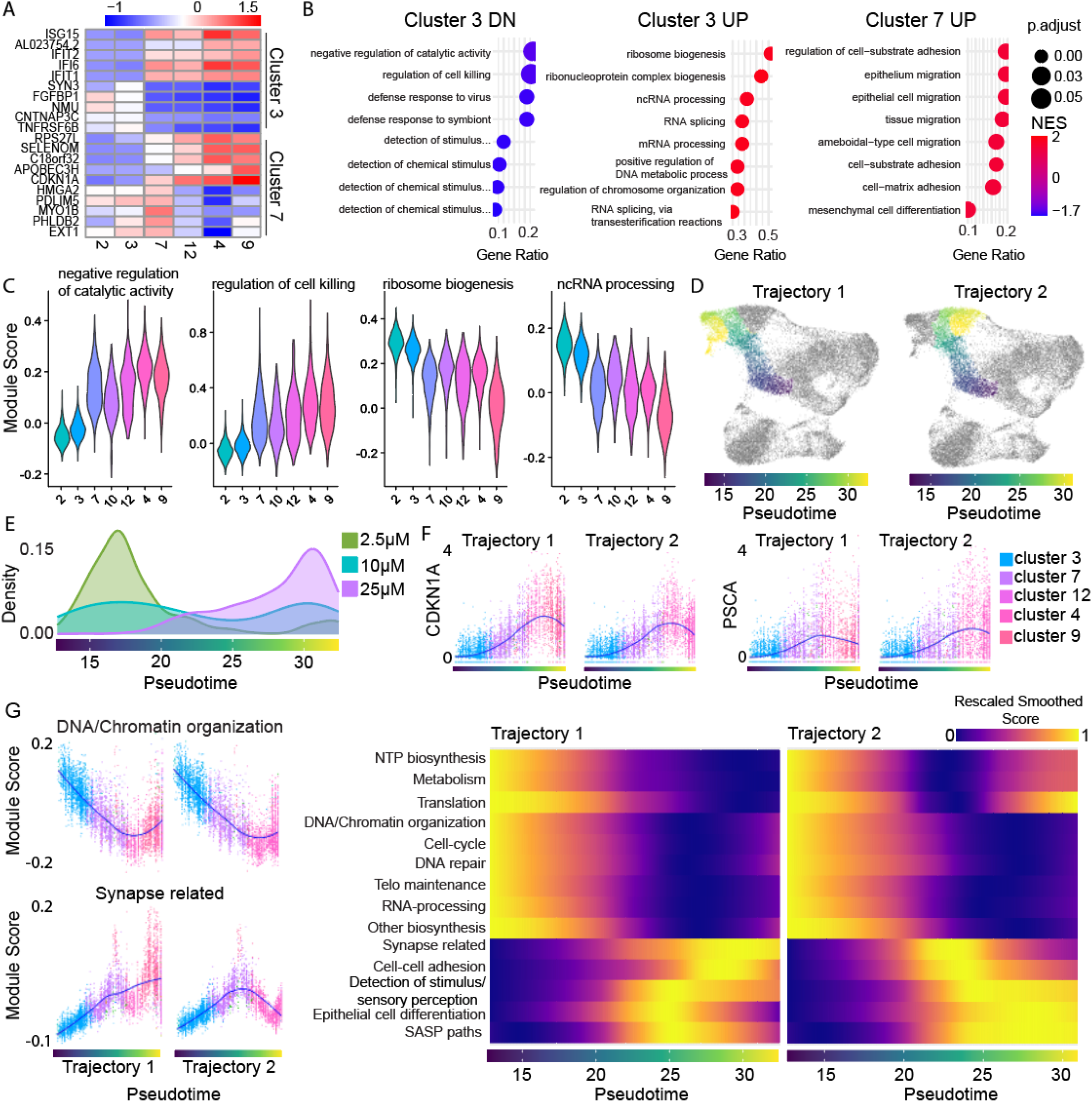
There is a continuous gradient of transcriptomic states along the dominant path to senescence. **(A)** Top 5 up- and down-regulated genes for cluster 3 (shallow quiescence) vs senescent cells (clusters 4, 9, 12) and cluster 7 (quiescence-senescence transition) vs senescent cells. **(B)** GSEA pathway analysis for results from DGE analysis in A. Top 8 pathways (ranked by p.adjusted) are shown. Cluster 7 had no significantly downregulated pathways. For full pathway names see Supplementary table 1. **(C)** Distribution of module scores by cluster for leading-edge genes in two selected up- and downregulated pathways for quiescent cluster 3 vs. senescent cells. **(D)** UMAP with cells colored by progress along pseudotime trajectory. Trajectory branches visualized separately as “Trajectory 1” and “Trajectory 2”. **(E)** Etoposide dose distribution of cells over pseudotime. **(F)** Expression of select top senescent markers over pseudotime. Cells colored by cluster. Loess fit visualized in blue. **(G) Left:** Module scores (see methods) for grouped biological processes over pseudotime. Scores for one select up- and down-regulated pathway group over pseudotime shown as single-cell data. Cells colored by cluster. Loess fit in blue. **Right:** Heatmap of all pathway groups over pseudotime. Color represents the Loess fit value for each module score vs. pseudotime scatter plot for the pathways listed along the y-axis. Module scores were rescaled for visualization on the same scale.

Within the quiescence-senescence transition cluster 7, some cells may more readily re-enter the cell cycle when compared to other senescent clusters based on the etoposide dose distribution in this cluster. Interestingly, pathways upregulated in cluster 7 relative to senescent clusters include “epithelium migration”, “tissue migration”, and cell-adhesion related pathways (Fig 4B and S4C), pathways that have been shown to promote metastasis and EMT in dormant cancer cells^31–33^. If such cells were to re-enter the cell cycle, they could pose a unique threat in a cancer context.

Clustering analysis defines clusters and “pseudobulks” the cells, sometimes masking heterogeneity within clusters. We scored cells based on their expression of leading-edge genes (see methods) for pathways differentially expressed in quiescent vs. senescent cells and found significant overlap in the distribution of scores for top pathways between adjacent clusters (Fig. 4C). We reasoned that this overlap between clusters may be due to continuous, graded changes in gene expression between cells on one “side” of the cluster compared to the other.

To test this idea, we performed pseudotime trajectory analysis using Monocle3^34,35^ (see methods), a computational method used to infer the progression of single cells through a biological process. Our goal was to identify transcriptional changes that occur as a function of progress through G0/G1-labelled clusters. We therefore subsetted the G0/G1 etoposide-treated cells and selected the cells closest to proliferating clusters as our ‘root’. Cells were then ordered and colored based on their progression along the learned trajectory (Fig 4D and S4D). Some cells with a G0/G1 label are in G1 phase of the cell cycle instead of arrested, and thus by including all G0/G1 cells, the resulting pseudotime trajectory spans all depths of quiescence into our most senescent cells. The trajectory branched partway, resulting in two different regions of “late pseudotime”, which were split into two trajectories for clarity during analysis (Fig 4D). We measured the distribution of each etoposide dose along pseudotime (Fig 4E). Progress along pseudotime correlates with a decreasing proportion of 2.5µM and an increasing proportion of 25µM cells. Given the changing proportion of doses along this trajectory, progress along pseudotime measures a decreasing probability of cell-cycle entry.

We examined if the top DE genes from our previous analyses were graded along pseudotime (Fig 4F and S4E). Indeed, CDKN1A and PSCA showed a graded increase over pseudotime, dipping slightly in cells that had progressed the farthest (Fig 4F). To extend this analysis beyond individual genes, we identified senescence-associated pathways in our data by selecting the significantly up- and down-regulated pathways for cluster 4 and cluster 9. We then used the leading-edge genes (most DE, see methods) from each of these pathways to calculate an expression score in every cell. Once the score was calculated, we plotted the score vs. pseudotime and performed a Loess fit to find the average score for each pathway across pseudotime. The Loess fit values for all pathways were then rescaled between 0 and 1 for visualization on the same color scale (S4F bottom).

Since many pathways were changed in our senescent cells, we combined pathways from the same general cellular functions as in Fig 3 such that the score was based on the leading-edge genes from all pathways grouped in that general cellular function (Fig 4G left and S4F top). To visualize all grouped pathways together over pseudotime, we generated a heatmap where the color corresponds to the fit and rescaled score (Fig 4G right). All downregulated pathways gradually declined over pseudotime, with NTP biosynthesis, metabolism, and translation-related pathways going back up in late pseudotime along Trajectory 2 but not Trajectory 1. The upregulated pathways showed the reverse behavior, with gradual increases across pseudotime. The varied behavior in later pseudotime reflects the heterogeneity of the expression of these phenotypes within the different senescent types. These results show that senescent phenotypes already begin to manifest early in pseudotime in cells with potential to re-enter the cell cycle. Further, quiescence represents a transitional state where cells exhibit expression signatures between that of proliferating and senescent cells.

### Protein biosynthesis rate is a key difference between quiescent and senescent cells

An outstanding question is how senescent cells permanently lose the ability to re-enter the cell cycle. We found that pathways associated with RNA-processing and translation comprised the two largest pathway groups downregulated in senescent clusters, after cell-cycle related pathways (Fig 3B). These pathways were also the most downregulated in senescent cells when compared to quiescent cells directly (Fig 4B-C). RNA-processing and translation are essential for efficient protein biosynthesis, which is in turn essential for the cell cycle, particularly in cancer cells^36,37^. Therefore, a decrease in RNA-processing and translation can impinge directly on the cell cycle.

We hypothesized that a decrease in protein biosynthesis might be predictive of an inability to re-enter the cell cycle. To test this, we treated MCF10A cells with 2.5µM, 10µM, or 25µM etoposide for 24h, followed by a 6-day drug-free recovery. At the end of the 6d, we treated cells for 30 min with O-propargyl-puromycin (OPP), which incorporates into actively translating proteins and measures global translation rate at the single-cell level. We detected a dose-dependent decrease in average and integrated OPP intensity (Fig 5A and S5B, respectively) in senescent cells, indicating reduced protein biosynthesis.

**Figure 5.**
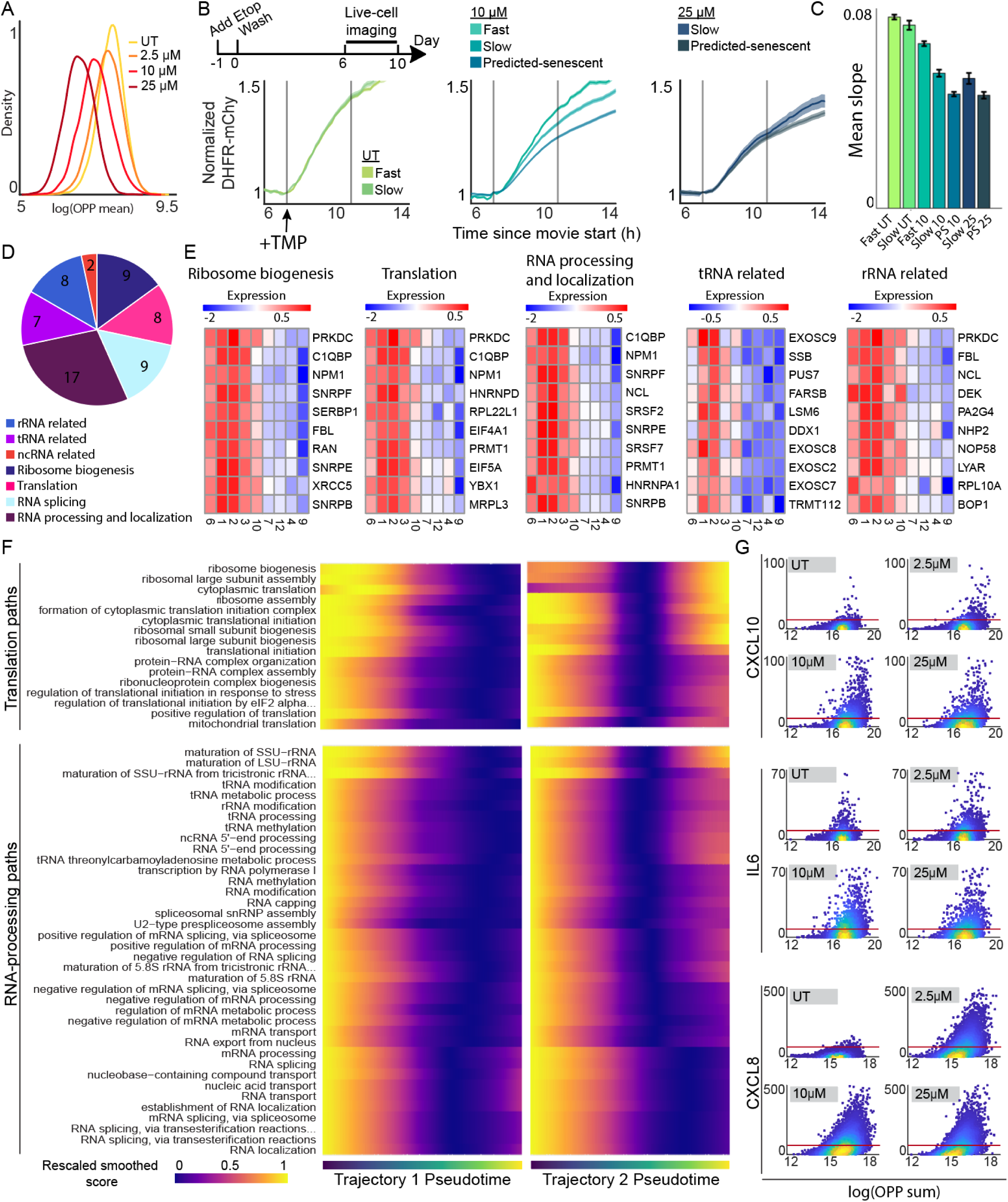
Protein biosynthesis declines along the quiescence-senescence continuum. **(A)** MCF10A cells were treated with 2.5 μM, 10μM, or 25μM etoposide for 24h followed by a 6d drug-free recovery. Cells were treated with OPP for 30 min prior to fixation. Each cell’s OPP intensity was normalized to the cell area and plotted as histograms for each etoposide dose. **(B)** Experimental timeline (top). MCF10A cells expressing the CDK2 activity reporter and a DHFR-mCherry translation reporter were imaged by live-cell microscopy for 96h from 6d-10d after etoposide washout. TMP was added at the first gray bar to induce the DHFR-mCherry reporter, and the mCherry signal (each cell normalized to its average signal before TMP) was plotted for fast-cycling, slow-cycling, or predicted-senescent cells for each dose of etoposide. Dark line is the average for each group, shading is the 95% confidence interval. Grey bars bracket the frames used to calculate the mCherry signal slope plotted in C. **(C)** A line was fit to the initial slope of mCherry accumulation after TMP addition. The average slope for each cell-cycle behavior group for each dose is shown. PS is predicted-senescent. **(D)** All significantly downregulated pathways from the RNA-processing and translation pathway groups in Fig. 3C for senescent clusters 4 and 9 were broken down into more specific pathway groups. **(E)** Average expression for select leading-edge genes by cluster from groups of pathways in D. **(F)** Heatmap of all GO biological processes from the RNA-processing and translation pathway groups over pseudotime. Colormap represents the Loess fit value for each module score over pseudotime. Module scores were rescaled for visualization on the same scale. For full pathway names see Supplementary table 1. **(G)** Cells were treated with 2.5 μM, 10μM, or 25μM etoposide for 24h followed by a 6d drug-free recovery. OPP was multiplexed with RNA FISH for three SASP genes and imaged. RNA FISH puncta number and total OPP per cell were quantified and plotted as a single-cell density scatter. Red line is drawn at the 95^th^ percentile of untreated cells.

To determine if translation rate could predict if cells were quiescent or senescent, we used a dihydrofolate reductase (DHFR)-trimethoprim (TMP) protein stabilization system fused to mCherry^19^, where the rate of mCherry accumulation upon TMP treatment reflects protein biosynthesis rate (S5C). We treated cells with 10µM or 25µM etoposide for 24h and then tracked them from 6-10d post-recovery (Fig 5B top) to measure CDK2 activity and mCherry accumulation rate (Fig 5B and S5D-E). We found that slow-cycling cells had a higher biosynthesis rate than non-cycling cells, in a dose-dependent manner (Fig 5B-C and S5F), identifying reduced biosynthesis as a distinguishing factor between quiescent and senescent cells.

### Protein biosynthesis is impaired in senescence, particularly in cluster 9

Protein biosynthesis relies on many important biological processes upstream. To understand which steps of this process may be driving a decrease in protein biosynthesis following senescence induction, we examined the DE pathways in our senescent clusters identified in Fig 3. We further broke down the translation and RNA-processing related pathways for clusters 4, 9, 10, and 7 by similarity (Fig. 5D, Supplementary table 1). We noted downregulation of biological processes that could affect protein biosynthesis at multiple steps, including rRNA related pathways, tRNA related pathways, and ribosome biogenesis and assembly (Fig. 5E). Examples of the strongly down-regulated genes include NPM1, EIF4A1, and exosome complex components. The largest number of pathways came from general RNA-processing and localization.

These data suggest that protein biosynthesis is decreased in senescent cells due to a multi-step failure, where no one step is solely responsible. In addition to the direct changes in RNA-processing, ribosome biogenesis, and translation initiation, we also measured a decrease in NTP biosynthesis and metabolism related pathways (Fig. 4G and S5I) in cells farther along pseudotime. Knockout of key regulators of NTP biosynthesis can cause cell-cycle arrest^38^, and NTP biosynthesis is coupled with protein biosynthesis in some cancers^39^. Thus, changes in these processes could indirectly affect protein biosynthesis and cell-cycle progression as well.

### Cells that express SASP factors have higher protein biosynthesis than cells that do not

The decrease in protein biosynthesis is interesting because translation has been shown to be especially important for maintaining the SASP^40–42^. We knew from our clustering and pseudotime analysis that pseudotime Trajectory 2 ended in cells that showed greater expression of SASP pathways than Trajectory 1. To test whether Trajectory 2 retained expression of pathways associated with translation and RNA processing, we used the same analysis strategy as in Fig 4, plotting the expression score of leading-edge genes over pseudotime, rescaled between 0 and 1. However, here we plotted this score for every individual GO biological process associated with translation or RNA processing (Fig 5F and S5G). We found that a majority of pathways associated with ribosome biogenesis, ribosome assembly, and translation initiation increased expression in late pseudotime along Trajectory 2 but not Trajectory 1, as we hypothesized.

To experimentally test this phenotype in cells, we multiplexed RNA FISH for three SASP factors, IL6, IL8, or CXCL10 with the OPP translation assay in the same single cells that had been treated with 2.5µM, 10µM, or 25µM etoposide 6d prior (Fig 5G). We found a dose-dependent increase in expression of all three SASP factors. In agreement with our scRNA -seq results, only a subset of the cells in each sample showed increased expression of the SASP factors, even at 25µM where nearly all cells are senescent. This SASP expression was highest in the cells with the greatest translation rate as measured by OPP. Expression of the SASP can be detrimental if cells are not properly cleared^43,44^. Therefore, future work investigating the pathways that are expressed differentially between Trajectory 1 and Trajectory 2 may offer new insights into suppressing the SASP without suppressing cell-cycle arrest.

### Expression of the SASP is higher in cells that have undergone a mitotic slip

Because the SASP can alter the surrounding tissue microenvironment and is thought to be a driver of age-related pathologies^43,44^, there is significant interest in characterizing the SASP at the single cell level. It was surprising that SASP expression was variable across our senescent clusters, with cluster 4 containing the most upregulated pathways associated with the SASP (Fig. 3B). We plotted the score for leading-edge SASP-related genes from our data set over pseudotime (Fig.6A) and saw that Trajectory 2, which is connected to the cluster of cells undergoing mitotic slippage, showed a graded and steady increase in expression. By contrast, Trajectory 1 showed an initial increase followed by a decrease in SASP-related genes, in agreement with our cluster analysis. We also used an existing SASP gene list from the Molecular Signatures Database (MSigDB)^29,45^ and scored cells based on their expression of these SASP factors (Fig. 6B). When visualizing the existing SASP gene list on our UMAP, we saw that cells near cluster 4 and cluster 10, the mitotic slip cluster, scored the highest for SASP expression.

**Figure 6.**
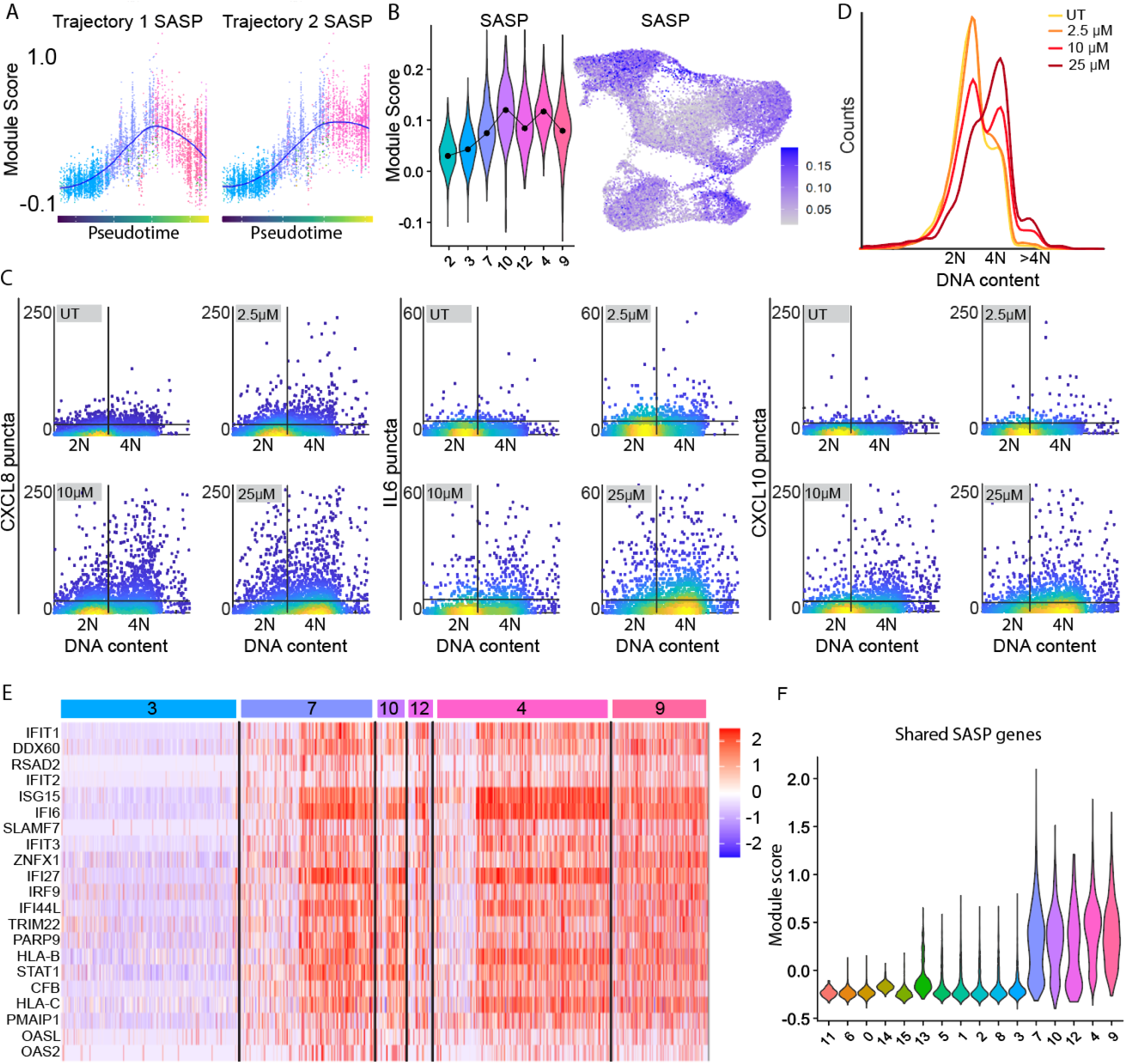
The SASP is expressed heterogeneously in senescent cells. **(A)** Module score for leading-edge genes from SASP-related biological processes over pseudotime. Loess fit in blue. Colored by cluster as in Fig. 4F. **(B) Left:** Distributions of module scores for REACTOME_SENESCENCE_ASSOCIATED_SECRETORY_PHENOTYPE_SASP gene set gene set by cluster. **Right:** UMAP colored by module score forSASP gene set on the left. **(C)** Cells were left untreated (UT) or treated with 10μM etoposide (2.5μM and 25μM in Fig. S6A) for 24h before being washed and allowed to recover for 6 d. Number of RNA FISH puncta for three SASP genes vs. DNA content is plotted as a single-cell density scatter. Horizontal line is the 95^th^ percentile of untreated puncta for each SASP factor. Vertical line is the saddle point between 2N and 4N DNA content (measured by Hoechst dye). **(D)** Distribution of DNA content for each dose. **(E)** Single-cell expression levels for SASP genes expressed in all senescent clusters. Quiescent cluster 3 and quiescence-senescence-transition cluster 7 shown for contrast. **(F)** Module scores for the same genes shown in E, by cluster. Clusters labeled at the bottom.

To test experimentally whether cells that arrest via a mitotic slip are the senescent cells with highest SASP expression, we performed RNA FISH for IL6, IL8, and CXCL10. We found that cells with 4N DNA content, many or most of which have undergone a mitotic slip at 10 µM and 25 µM (Fig. 2D), had the highest expression of SASP RNAs (Fig. 6C-D and S6A). While cells arresting by the G0/G1 trajectory had upregulation of some SASP-related pathways (Fig. 3, cluster 7), these pathways were related more to wound-healing and extracellular matrix organization rather than the defense-response related pathways that we saw for cells that had undergone a mitotic slip (Fig. S6B and S6C). These data suggest that not only are there different senescent types in our population, but the type of senescence that arises in each cell is tied, at least in part, to the initial mode of arrest that cell underwent.

### A subset of SASP genes is expressed in all types of senescence

Because different clusters showed differing expression of SASP genes, we tested whether a subset of SASP genes was expressed in all senescent clusters. We found 21 SASP genes that were universally upregulated (see methods) in every senescent cluster at every dose of etoposide (Fig. 6E). Quiescent cluster 3 was used for contrast. We plotted the expression levels of these SASP factors in single cells and confirmed that all senescent clusters showed expression of these factors (Fig. 6F). Interestingly, many of these specific genes are double-strand (ds) RNA-sensing proteins (OASL, OAS2) or interferon-inducible (IFI) antiviral genes (IFIT1, IFIT2, IFI6) downstream of these RNA-sensing proteins. While the upstream stresses driving the SASP can be varied, these data suggest that dsRNA accumulation may be a common stress driving SASP factor expression across senescent types.

### A gene set to identify TIS cells

The lack of clear biomarkers to distinguish between quiescent and senescent cells is a persistent problem in the field. We generated a gene set based on genes represented in all senescent clusters that did not overlap with quiescent cells (see methods, Fig. S7A) and scored cells based on their expression of this gene set (Fig. 7A and S7A). Some of the top markers for senescent cells showed elevated expression in untreated quiescent cells, and we therefore removed them to generate a senescence-specific list of genes. This analysis revealed multiple genes that were shared across the senotypes. Importantly, the expression of genes in this list was not dose-dependent or driven by a changing proportion of 25µM cells, since it was able to identify senescent cells across all doses (Fig S7B-C). The genes in our gene set were generally highly expressed in senescent clusters, showed low or no expression in other clusters, and included several new senescent biomarkers, including IFI6, IFI27, SAMD9L, RSAD2, RNASE7, SPINK5/6, SELENOM, IFIT3, OAS2, and HEPH (Fig 7B).

**Figure 7.**
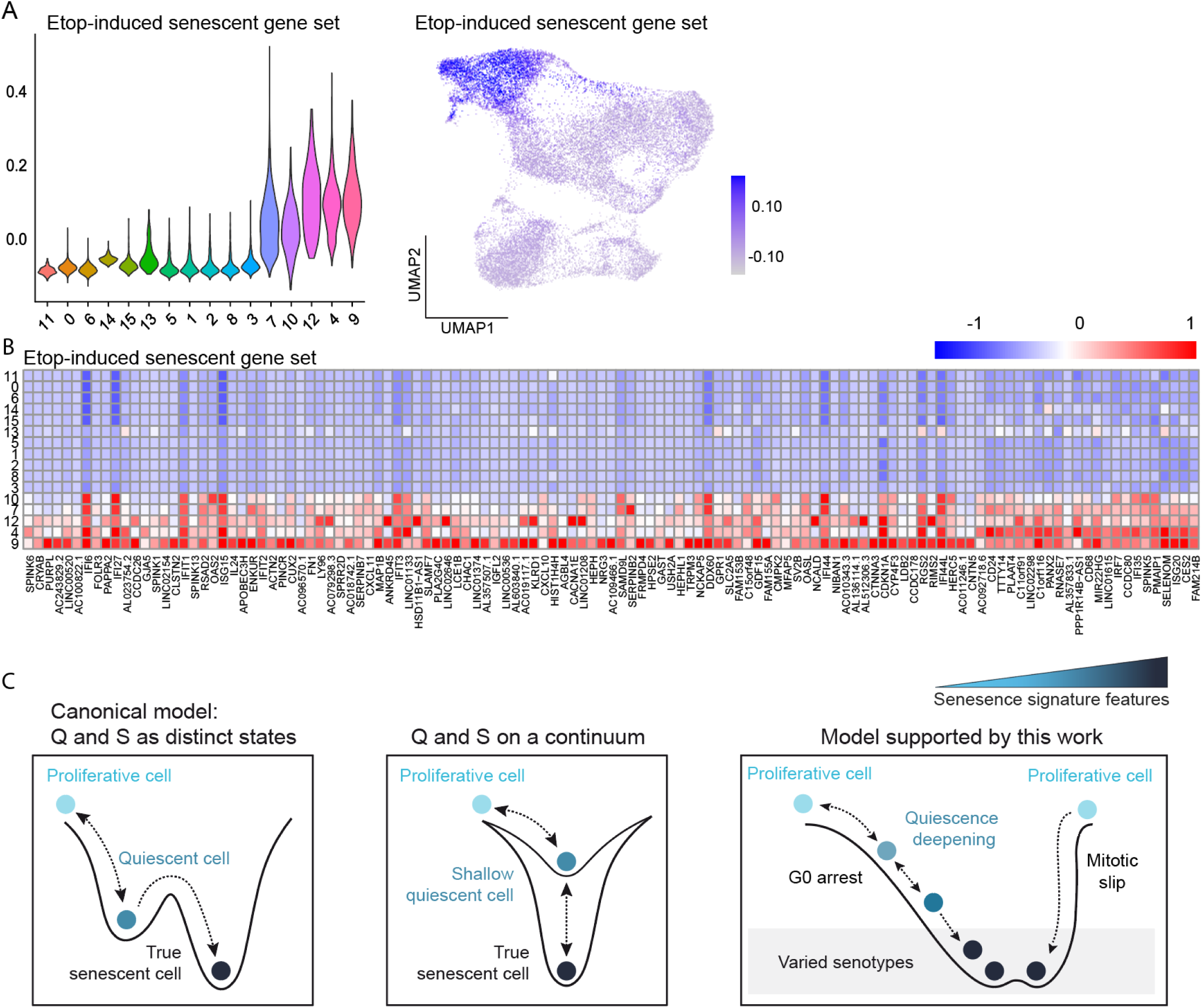
A new senescent gene set to identify diverse senescent cell types following etoposide. **(A)** Distribution of module scores for cells from each cluster based on our senescent gene set (left). UMAP with dots colored by module score for our senescent gene set (right). **(B)** Average expression of all genes in our senescent gene set by cluster. **(C) Left:** Canonical model of a switch-like transition between quiescence and senescence. **Middle:** Simple continuum model of cell-cycle withdrawal where quiescence and senescence represent different levels on the same continuum of withdrawal. **Right:** Model of cell-cycle arrest supported by this work. In the dominant path, cells complete mitosis and first enter a shallow quiescence (G0). In this path, quiescent cells represent a reversible intermediate between proliferation and senescence and show a graded expression of senescence genes. In a rarer path, cells skip mitosis and arrest with 4N DNA content. These cells transit directly to a senescent state, bypassing the quiescence deepening observed along the dominant path.

## Discussion

The relationship between different quiescence and senescence remains incompletely understood (Fig. 7C), in large part due to the lack of specific and consistent biomarkers. Here, we paired scRNA-seq with single-cell time-lapse microscopy to identify ground-truth senescent cells based on a functional definition of senescence: the long-term lack of proliferation. Our data revealed a number of findings that we discuss below.

Related to the cell cycle itself, we found that rapidly cycling cells after etoposide treatment have a transcriptionally encoded “memory” from the initial etoposide stress, even after 6d. Further work characterizing the gene expression signatures in cells that resume cycling after chemotherapy treatment relative to untreated cycling cells could expose vulnerabilities to eliminate this population. We also found two distinct arrest paths, or bridges, from the cell cycle to senescence, a dominant mitosis-to-G0 path and a less common mitotic slip path. While cells taking the first path progress from G0/quiescence to senescence in a graded manner, cells arresting by the mitotic slip path appear to move directly from cycling to senescence based on the fact that they move straight from the G2/M cluster to the senescent clusters (Fig. 7C). This mitotic slip path was primarily occupied by 10 and 25µM cells, suggesting that higher levels of damage may drive cells towards this path over the mitosis-to-G0 arrest path. Cells that are in S phase when they are first treated with etoposide may also be more susceptible to this arrest path^46^.

Related to senescence, we identified several transcriptomic clusters of senescent cells, or senotypes. Thus, senescence heterogeneity arises not only from different treatments or cell types, as previously reported^47–50^, but also within the same population of cells undergoing the same treatment. Although these senotypes were clustered finitely for analysis and showed cluster-to-cluster differences, there was a continuous distribution of cells across the senescent region of our UMAP. This suggests that the gene expression distinctions between these senotypes are graded, with single senescent cell transcriptomes varying across the entire senescent transcriptomic space. Prior work, including ours, noted variability in senescence markers under identical conditions^16,51^. Our findings suggest that this variability reflects both incomplete senescence induction and the presence of senescent cells lacking expression of certain canonical markers.

Interestingly, these different senotypes arose following the two distinct arrest paths. Mitotic slipping cells expressed higher levels of inflammation and protein biosynthesis-related pathways compared to cells taking the mitosis-to-G0 path, which showed higher expression of pro-growth and EMT-related pathways such as ‘wound healing’. In the context of cancer, the pro-inflammatory SASP can recruit the immune system to initiate clearance, whereas the pro-growth arm of the SASP can drive surrounding cells to proliferate^7,8^. Further work is needed to understand how these different senotypes may affect surrounding cells. The SASP requires a basic level of cellular function for coordinated expression, synthesis, and secretion of the program. Cluster 9 cells experiencing a high level of stress – marked by high p21, low RNA processing/translation, metabolic stress, and distance from cycling etoposide-treated cells – likely lack this capacity. Indeed, cluster 9 cells showed reduced SASP-associated pathways, whereas SASP-expressing cells in cluster 4 had higher expression of genes linked to translation, RNA processing, NTP biosynthesis, and metabolism.

We identify changes in protein synthesis as a key difference between quiescence and senescence. It has been shown that changes in RNA processing and translation are a part of the DNA damage response^52–54^ and reduced mTOR and translation signaling are also associated with senescence^55,56^. Here we reveal that these gene changes span a range of processes involved in the processing of mRNA, tRNA, rRNA, and lncRNAs, ribosome biogenesis and assembly, and direct translational control. Importantly, the differential expression of these pathways between quiescent and senescent cells places renewed importance on protein biosynthesis as a uniquely useful predictor of future cell-cycle re-entry, with lack of protein biosynthesis as a causal determinant of irreversible cell-cycle withdrawal.

We defined a core gene set shared across all senescent clusters, distinguishing them from non-senescent clusters, along with a subset of upregulated SASP genes that are expressed in all of our senescent cells. These gene sets may enable transcriptional-level identification for sequencing applications, and thus should be validated in other cell types and in other modes of senescence induction. Future studies will be needed to elucidate the mechanistic role, if any, of key markers in senescent cell biology.

Lastly, we found that our senescent population was connected to the treated, cycling cells by a continuous UMAP “bridge”. Using pseudotime trajectory analysis of the mitosis-to-G0 path, we found that rather than a switch like change, multiple senescent phenotypes begin to manifest early and gradually, even in shallow quiescent cells. Thus, in the context studied here, quiescence represents a path by which cells move from proliferation to senescence, and cells positioned along this continuum express gradually varying levels of senescence markers.

This model represents an emerging view of the relationship between quiescence and senescence, where quiescence and senescence are not viewed as binary distinct states, but instead exist on a continuum with declining probabilities of cell-cycle re-entry (Fig. 7C). This model is consistent with previous work from our lab examining eight protein-based senescence biomarkers^16^. However, it was surprising to see that the graded model of cell-cycle arrest extends across every senescence pathway we identified here in our ground-truth senescent cells, providing transcriptome-wide support for this model. A handful of other studies also support the continuum model. One study that used bulk RNA-seq found that prolonged serum starvation of fibroblasts, a reversible quiescence-inducing treatment, led to some transcriptional changes that are typical of senescence at the bulk level, such as lysosome biogenesis and autophagy^57^. While we also observed changes in some of these genes, they were less pronounced, suggesting that arrested cells may lie on the quiescence-senescence continuum, but the rate at which different senescent phenotypes emerge is likely context dependent. Moreover, not all senescent phenotypes may change in every arrested condition, warranting further study to understand their overlap and trajectories. Another study used multi-plexed imaging to examine arrest induced by serum starvation, etoposide, and oxidative stress^51^, concluding that senescence represents a G1-like arrested state that eventually converges on a single senescent state following different arrest trajectories. This work was limited to primarily cell cycle-related proteins and a few canonical senescent markers, explaining why the data appeared to converge on a single senescent phenotype. In our study here, with measurements across the entire transcriptome in ground-truth senescent cells, we found that multiple senescent phenotypes arose. Another recent multi-omics study of passage limited fibroblasts^58^ found that senescent-like changes manifest early, before replicative senescence was reached, suggesting this model likely extends to replicative aging as well. Future work delineating the specific genes and processes that are universally changed in irreversibly arrested cells is warranted.

At what point along this continuum of withdrawal do cells become irreversibly arrested and why? We found that irreversibly arrested cells showed perturbations in multiple genes and pathways that likely impair cell-cycle entry and progression in multiple ways. This suggests that no single, switch-like expression change drives the irreversibility of cell-cycle withdrawal. Instead, as cells progress along the continuum, multiple different senescent traits that deepen arrest intensify, creating a greater and greater barrier to cell-cycle re-entry, which eventually becomes insurmountable.

## Methods

### Cell lines and culture media

MCF10A (ATCC CRL−10317) cells were obtained from ATCC and grown in DMEM/F12 supplemented with 5% horse serum, 20 ng/ml EGF, 10 μg/ml insulin, 0.5 μg/ml hydrocortisone, 100 ng/ml cholera toxin, and 100 μg/mL of penicillin and streptomycin. Cells were imaged in phenol red-free full growth media during live-cell movies. Cells were cultured and imaged in a humidified incubator at 5% CO_2_ and 37 °C.

### Drug treatments

MCF10A cells were split 1:10 (from a confluent plate) into a plastic 10cm culture plate before being treated with 2.5, 10, or 25μM etoposide the following day for 24 h. Cells were then washed twice with PBS before being returned to full-growth media. Full-growth media was refreshed every 3d during the drug recovery period. 24 h prior to imaging, the etoposide-released cells were trypsinized and replated onto a collagen coated (1:50 dilution in water) (Advanced BioMatrix, No. 5005) 96-well glass-bottom plate (Cellvis Cat. No. P96−1.5H-N). Cells were plated at 1000, 2000, 3000, and 4000, cells per well for untreated, 2.5, 10, or 25μM etoposide, respectively. Cells were plated at 3000 cells per well for immunofluorescent imaging.

### Immunofluorescence imaging

Cells were seeded onto a collagen coated (1:50 dilution in water) (Advanced BioMatrix, No. 5005) 96-well glass-bottom plate (Cellvis Cat. No. P96-1.5H-N) 24 h prior to fixation. Cells were fixed for 15 minutes with 4% PFA in PBS then permeabilized at room temperature with 0.1% TritonX for 15 min. Cells were then washed with PBS and blocked with 3% Bovine Serum Albumin (BSA) for 1 h at room temperature. Primary antibodies were incubated overnight in 3% BSA at 4 °C and secondary antibodies were incubated for 1-2 h in 3% BSA at room temperature. Nuclei were labelled with Hoechst at 1:10,000 in PBS for 10 min. The whole cell was stained with succinimidyl ester 488 at 1:10,000 in PBS for 30 minutes. Cells were washed with 100 μL per well of PBS between every step. All images were obtained using a 10X, 0.4 numerical aperture objective on a Nikon TiE microscope. To visualize IL8, we blocked secretion for 6h with 5 µg/mL Brefeldin A before fixing and staining. For OPP translation assay, cells were treated according to the manufacturer’s protocol (Click-iT Plus OPP Alexa Fluor 488 Protein Synthesis Assay Kit, ThermoFisher C10456), with cells incubated with OPP reaction component A for 30 minutes prior to fixation.

### RNA FISH imaging

Cells were seeded onto a collagen coated (1:50 dilution in water) (Advanced BioMatrix, No. 5005) 96-well glass-bottom plate (Cellvis Cat. No. P96-1.5H-N) 24 h prior to fixation. Cells were fixed for 15 minutes with 4% PFA in PBS. IL6 (VA6−12712-VC), IL8 (VA6-13192-VC), and CXCL10 (VA6-13729-VC) mRNA were visualized according to the manufacturer’s protocol (ViewRNA ISH Cell Assay Kit, ThermoFisher QVC0001), with cells permeabilized for 30 min and mRNA probes hybridized for 4h at 40C.

### Time-lapse microscopy

Cells were plated 24h prior to imaging and full-growth media was replaced with phenol red-free full-growth media. Images were taken for each fluorescent channel every 12 min at two sites per well that were spaced 2 mm apart. Total exposure across all fluorescent channels was kept below 800 ms. Cells were imaged in a humidified, 37 °C chamber at 5% CO_2_. All images were obtained using a 10X, 0.4 numerical aperture objective on a Nikon TiE microscope. The tracking code is available for download here: https://github.com/scappell/Cell_tracking.

### Image processing and quantification

Image processing and cell tracking were performed using MATLAB Mathworks 2017b as previously described^59^. Quantification of 53BP1 puncta was measured as previously described^60^. Nuclear signals (phospho-Rb, 53BP1, p21, and Lamin B1) were quantified from a nuclear mask (median nuclear intensity). Cytoplasmic signals (LAMP1 and IL8) were quantified from a cytoplasmic mask (median cytoplasmic intensity) derived from succinimidyl ester total protein stain. RNA FISH puncta detection was performed using Revvity’s Harmony High-Content Imaging and Analysis Software. All puncta inside the cytoplasmic mask belonging to each nucleus were counted.

### Antibodies and reagents

Antibodies against pRb (S807/811) D20B12 XP (8516), LAMP1 D2D11 XP (9091), p21 Waf1/Cip1 (12D1) (2947), and Lamin B1 (D9V6H) (13435) were purchased from CST and used at 1:500, 1:1000, 1:250, and 1:1000 dilutions, respectively. Antibodies against 53BP1 (612523), and IL-8 (550419) were purchased from BD and were all used at dilutions of 1:1000. All secondary antibodies, Goat anti-Mouse IgG (H + L) Cross-Adsorbed Secondary Antibody, Cyanine3 (A10521), Goat anti-Rabbit IgG (H + L) Cross-Adsorbed Secondary Antibody, Cyanine3 (A10520), Goat anti-Mouse IgG (H + L) Highly Cross-Adsorbed Secondary Antibody, Alexa Fluor 647 (A-21236), Goat anti-Rabbit IgG (H + L) Highly Cross-Adsorbed Secondary Antibody, Alexa Fluor 647 (A-21245) were purchased from Thermo Scientific and used at 1:1000 dilutions. IL6 FISH mRNA probe set (VA6−12712-VC), IL8 FISH mRNA probe set (VA6-13192-VC), and CXCL10 mRNA probe set (VA6-13729-VC) were purchased from Thermo Scientific. The ViewRNA ISH Cell Assay Kit was purchased from Thermo Scientific (QVC0001). CF 488A succinimidyl ester (SCJ4600018) was purchased from Sigma and used at a 1:10,000 dilution. Hoechst 33342 was purchased from Thermo Scientific (H3570) and used at a 1:10,000 dilution. Etoposide (E1383), and Brefeldin A (B7651) were purchased from Sigma. Click-iT Plus OPP Alexa Fluor 488 Protein Synthesis Assay Kit (C10456) was purchased from Thermo Scientific.

### Single-Cell RNA Sequencing

Single-cell RNA sequencing (scRNA-seq) libraries were generated using the 10x Genomics Chromium platform with the 3’ mRNA capture kit. Single-cell suspensions were prepared following the 10x Genomics “Cell Preparation for Single Cell Protocols” guide with a viability of >95%. Single-cell encapsulation, barcoding, cDNA amplification, library preparation, and sequencing were performed by the University of Colorado Anschutz Genomics Shared Resource facility. Cells were encapsulated into droplets using the Chromium Controller, where reverse transcription occurred within individual gel beads-in-emulsion (GEMs). Following barcoding and cDNA amplification, sequencing libraries were constructed according to the manufacturer’s protocol. Libraries were sequenced on an Illumina NovaSeqX to a depth of 81,810 reads per cell for untreated, 74,214 reads per cell for 2.5µM, 81,254 reads per cell for 10µM, and 76,973 reads per cell for 25µM.

### scRNA-seq Data Processing

scRNA-seq data were processed and analyzed using the Seurat^21,22^ (v5.1.0) package in R (v4.4.1). Raw sequencing reads were pre-processed using CellRanger (v7.1.0) and aligned to the Human (GRCh38) 2020-A reference genome. Gene-cell count matrices were generated, and low-quality cells were filtered out based on the number of detected genes, unique molecular identifiers (UMIs), and mitochondrial gene percentage. Cells with fewer than 3,500 genes were excluded to remove low-quality cells. Cells with greater than 15% mitochondrial gene expression were also removed. Data were log-normalized using Seurat’s NormalizeData() function with a scale factor of 10,000.

### Feature Selection and Dimensionality Reduction

Highly variable genes (HVGs) were identified using FindVariableFeatures(), and data were scaled using ScaleData(). Principal component analysis (PCA) was performed using RunPCA() on the top N principal components (PCs). The optimal number of PCs (30) was determined using an elbow plot and JackStraw analysis^21,22^ and used in downstream dimension reduction.

### Clustering and Visualization

Cells were clustered using the shared nearest neighbor (SNN) graph-based clustering algorithm with FindNeighbors() and FindClusters() at a resolution of 0.8. Optimal clustering resolution was determined by iterative clustering and select downstream analysis at every resolution. Uniform Manifold Approximation and Projection (UMAP)^20^ was used for dimensionality reduction and visualization with RunUMAP(). Cell-cycle scoring was performed using Seurat’s CellCycleScoring() function, which assigns cell-cycle phase predictions based on predefined gene sets for G2/M and S phases. Any cells that aren’t categorized as G2/M or S are labeled G0/G1.

### Differential Expression Analysis

Differentially expressed genes (DEGs) between clusters were identified using FindMarkers() with the Wilcoxon rank-sum test. Multiple testing correction was performed using the Bonferroni method to adjust p-values. The expression of canonical markers was visualized using FeaturePlot(), VlnPlot(), and DotPlot(). Expression values shown in all single-gene UMAPs and heatmaps represent z-scored gene counts, which were normalized using NormalizeData() and subsequently scaled and centered with ScaleData().

### Gene Set Enrichment Analysis

Gene Set Enrichment Analysis (GSEA) was performed using the clusterProfiler package^61^ in R to identify enriched biological processes. The ranking metric was calculated as: Ranking Score=avg_log2FC×−log10(adj_pvalue), where avg_log2FC represents the average log2 fold change in gene expression, and adj_pvalue is the Benjamini-Hochberg adjusted p-value. This ranking method prioritizes genes that are both highly differentially expressed and statistically significant. Enrichment analysis was conducted using the gseGO() function, specifying the org.Hs.eg.db annotation package for human genes and the Biological Process (BP) ontology. The results were visualized using the gseaplot() function to display the leading enriched pathway and the dotplot() function to summarize the top enriched pathways. All analyses were performed in R using the following packages: clusterProfiler, org.Hs.eg.db, and ggplot2 for visualization.

### Identifying Shared SASP Genes

We first took the 118 leading edge genes for SASP-related pathways in our senescent clusters (clusters 4, 9, 10, 12) and then for each senescent cluster, we selected the top 50 of the 118 original genes. We then found the overlap between these lists, resulting in 21 SASP genes that were universally upregulated in every senescent cluster at every dose of etoposide (Fig 6D).

### Identifying the Etoposide-Induced Senescent Gene Set

To make a gene set to identify etoposide-induced senescent cells we combined our senescent clusters (clusters 4, 9, 10, 12) and performed DE expression analysis on our combined senescent cells vs all other cells sequenced. We selected the top 100 genes (based on Ranking Score=avg_log2FC×−log10 (adj_pvalue)) and combined these with the top 50 genes from DE analysis of cluster 4 alone vs all other cells, since this cluster was less robustly identified. We then found the unique genes from this combined list. Finally, any genes from this list that were also elevated in untreated G0/G1 cells were removed.

### Expression Score Calculation

Gene module scores were computed using AddModuleScore() for predefined gene sets, and results were visualized with FeaturePlot() and DoHeatmap().

### Pseudotime Analysis

Pseudotime trajectory inference was performed using Monocle3^34,35^, with preprocessed Seurat data converted via as.cell_data_set(). Trajectories were learned with learn_graph() and cells were ordered along pseudotime using order_cells().

## Data Availability

All sequencing data and processed count matrices will be deposited in Gene Expression Omnibus (GEO) (NCBI) (https://www.ncbi.nlm.nih.gov/geo/) upon publication. The analysis scripts are available upon request.

## Acknowledgements

We thank current and past members of the Spencer Laboratory for general discussion and insight over the course of the work, and Theresa Nahreini and the cell culture facility for cell sorting (RRID:SCR_018988). The Aria Fusion FACS sorter is supported by NIH grant S10OD021601. The RNA FISH imaging was performed at the Biofrontiers Advanced Light Microscopy Core directed by Joe Dragavon (RRID: SCR_018302). The Revvity Opera Phenix is supported by NIH grant 1S10OD025072. Single-cell capture, library generation and sequencing were performed at the University of Colorado Anschutz Medical Campus Genomics Shared Resource (RRID: SCR_021984), which is supported by the Cancer Center Support Grant (P30CA046934). We thank Dr. James DeGregori for providing facilities for sample preparation at Anschutz. This work was supported by an NIH T32 GM142607 (to B.F.), an NIH F31 CA284877 (to B.F.), an NIH Director’s New Innovator Award 1DP2CA238330-01 (to S.L.S) and an R01 R01AG082942 (to S.L.S).

## Author contributions

B.F. and S.L.S. designed the research; H.A., V.P., and B.F. conducted the research; B.F. analyzed the data; B.F. and S.L.S. conceived the project; and B.F. and S.L.S. wrote the paper; S.L.S. supervised the project.

## Corresponding author

Correspondence to <u>Sabrina L. Spencer</u>.

