## Supplemental Figures for "Single-cell RNA sequencing reveals distinct senotypes and a quiescence-senescence continuum at the transcriptome level following chemotherapy"

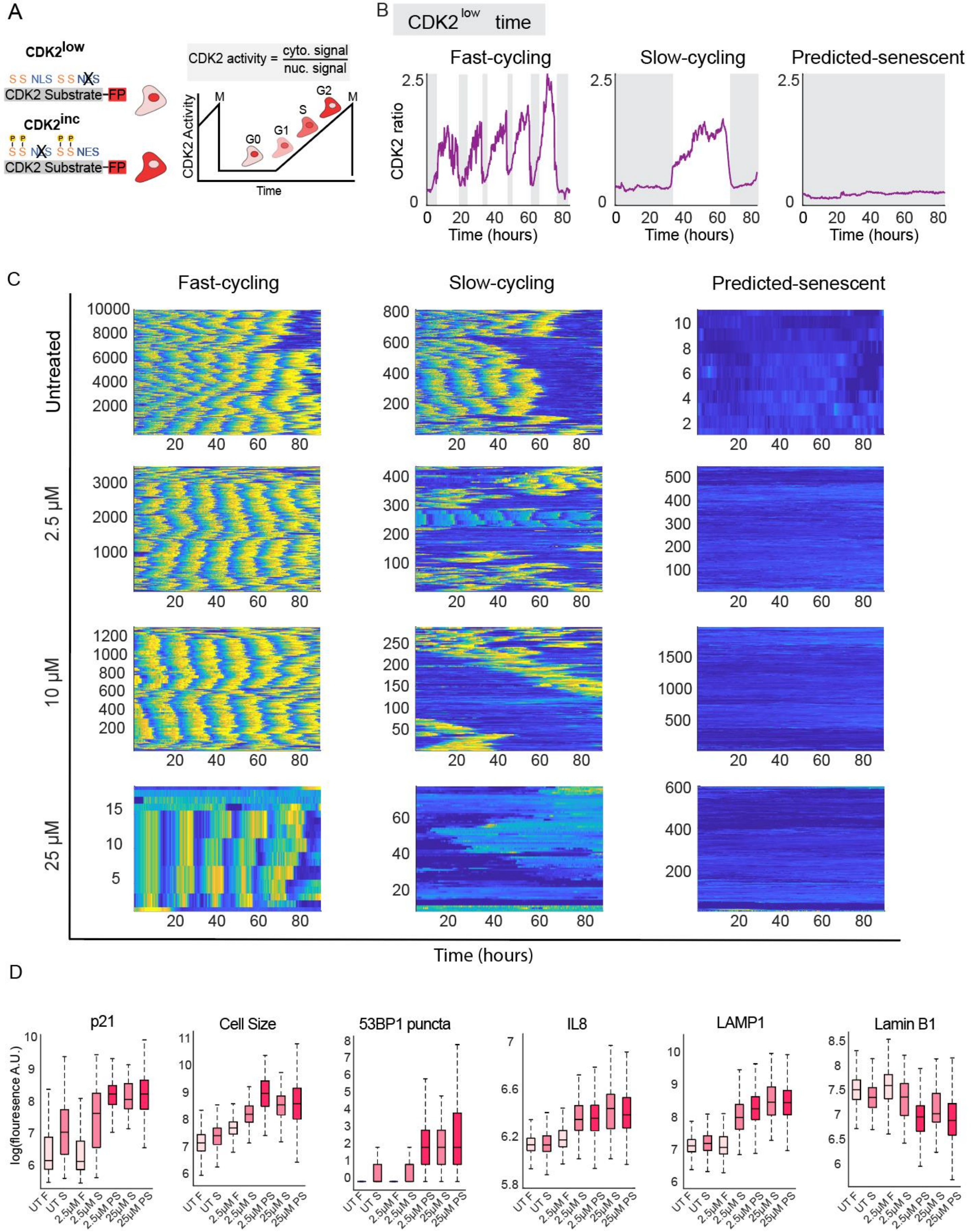

**Supplemental Figure 1. Time-lapse imaging of CDK2 activity resolves cell-cycle fates following etoposide. (A)** DHB-based CDK2 activity sensor first developed in Spencer et al.[19]. Sensor localizes to the nucleus when unphosphorylated. Phosphorylation by CDK2 leads to translocation of the sensor to the cytoplasm. NES, nuclear export signal. NLS, nuclear localization signal. FP, fluorescent protein. **(B)** MCF10A cells expressing the CDK2 activity sensor were treated with 2.5 μM, 10μM, or 25μM etoposide for 24h followed by a 6d drug-free recovery. Randomly selected single-cell CDK2-activity traces for fast-cycling, slow-cycling, and predicted-senescent cells at each dose. Grey shading corresponds to movie frames where the cell is not committed to cell cycle (CDK2 activity > 0.8). **(C)** Heatmaps of CDK2 activity (yellow: high, blue: low) as in Fig. 1 for cells in the fast-cycling, slow-cycling, and predicted-senescent categories for each etoposide dose. Cells are categorized based on cumulative $CDK2^{low}$ time. Cells with $CDK2^{low}$ time < 50h out of 96h total are categorized as fast-cycling. Cells that are continuously $CDK2^{low}$ for the entire movie are classified as predicted-senescent. **(D)** Cells from C were fixed and stained for canonical senescent biomarkers after the last movie frame. Cell-cycle histories were computationally linked to fixed-cell stains. Cells were categorized as fast-cycling, slow-cycling, or predicted-senescent based on time spent $CDK2^{low}$. These results show that etoposide dose modulates the proportion of predicted-senescent cells based on both CDK2 activity as well as their graded expression of canonical senescent markers as predicted by our previous work[16].

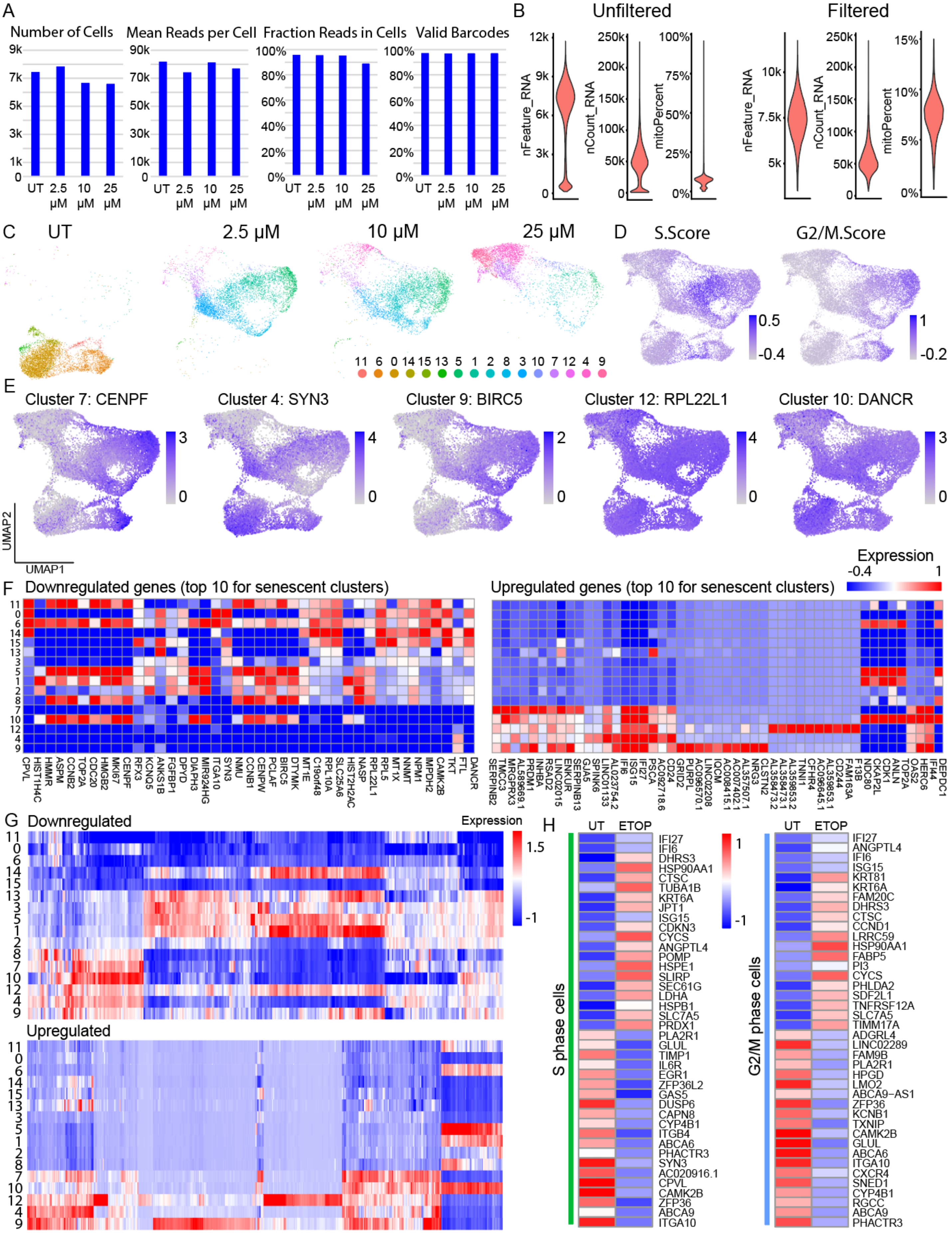
A
Number of Cells
Mean Reads per Cell
Fraction Reads in Cells
Valid Barcodes
UT
2.5 μM
10 μM
25 μM
B
Unfiltered
Filtered
nFeature_RNA
nCount_RNA
mitoPercent
C
UT
2.5 μM
10 μM
25 μM
11 6 0 14 15 13 5 1 2 8 3 10 7 12 4 9
D
S.Score
G2/M.Score
E
Cluster 7: CENPF
Cluster 4: SYN3
Cluster 9: BIRC5
Cluster 12: RPL22L1
Cluster 10: DANCR
UMAP1
UMAP2
F
Downregulated genes (top 10 for senescent clusters)
Upregulated genes (top 10 for senescent clusters)
Expression
G
Downregulated
Upregulated
Expression
H
UT
ETOP
S phase cells
G2/M phase cells
IFI27
IFI6
DHRS3
HSP90AA1
CTSC
TUBA1B
KRT6A
JPT1
ISG15
CDKN3
CYCS
ANGPTL4
POMP
HSPE1
SLIRP
SEC61G
LDHA
HSPB1
SLC7A5
PRDX1
PLA2R1
GLUL
TIMP1
IL6R
EGR1
ZFP36L2
GAS5
DUSP6
CAPN8
CYP4B1
ITGB4
ABCA6
PHACTR3
SYN3
AC020916.1
CPVL
CAMK2B
ZFP36
ABCA9
ITGA10
IFI27
ANGPTL4
IFI6
ISG15
KRT81
KRT6A
FAM20C
DHRS3
CTSC
CCND1
LRRC59
HSP90AA1
FABP5
PI3
CYCS
PHLDA2
SDF2L1
TNFRSF12A
SLC7A5
TIMM17A
ADGRL4
LINC02289
FAM9B
PLA2R1
HPGD
LMO2
ABCA9-AS1
ZFP36
KCNB1
TXNIP
CAMK2B
GLUL
ABCA6
ITGA10
CXCR4
SNED1
CYP4B1
RGCC
ABCA9
PHACTR3

**Supplemental Figure 2. scRNA-sequencing of etoposide treated cells. (A)** Quality control statistics for single-cell RNA sequencing data broken down by dose. **(B)** Feature count (nFeature_RNA), RNA molecule count (nCount_RNA), and percent of mitochondrial genes (mitoPercent) before (left) and after (right) filtering. Violin plots on the right correspond to cells used for analysis. **(C)** UMAP visualization of single cells for each etoposide dose. Color corresponds to the clusters in Fig. 2B. **(D)** Seurat cell-cycle scoring for S and G2/M cells. Color corresponds to relative module score where bluer means higher. **(E)** One top down-regulated gene for each senescent cluster. Color corresponds to normalized and scaled expression. **(F)** 10 most downregulated (left) and upregulated (right) genes for each senescent cluster were combined, and average expression of each gene was calculated (color) for each cluster. **(G)** 50 most downregulated (left) and upregulated (right) genes for each senescent cluster were combined and average expression of each gene was calculated (color) for each cluster. **(H)** Average expression of top 20 up-regulated and down-regulated genes between untreated and etoposide-treated S phase cells (left) and G2/M phase cells (right).

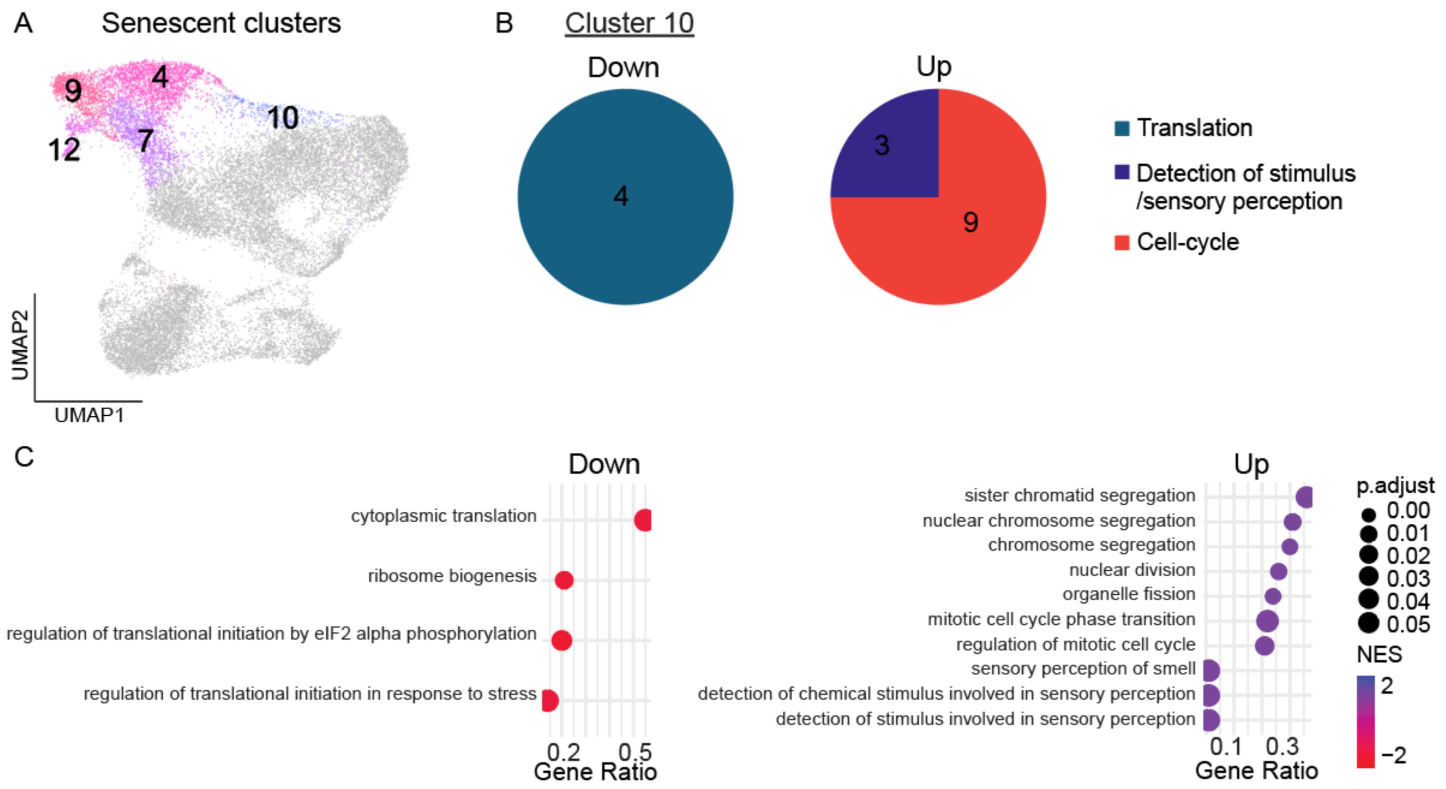


**Supplemental Figure 3. Pathway analysis of cluster 10 senescent cells. (A)** UMAP visualization of senescent clusters. **(B)** Significant up- and down-regulated pathways for cluster 10 relative to all other cells were grouped by general biological process (see methods, Supplementary table 1). These cells are senescent since this cluster is made up of 25μM cells that are not cycling based on time-lapse imaging. However, they are (incorrectly) predicted to be G2/M by Seurat due to upregulation of mitotic pathways, likely a result of having undergone a mitotic slip. **(C)** Gene set enrichment analysis (GSEA) for cluster 10. Top 10 pathways (by p.adjusted) up-regulated and all down-regulated pathways are shown. Gene ratio for each pathway (proportion of genes in each pathway that are significantly DE compared to the total number of genes in that pathway) plotted on the x-axis. Dots colored by the normalized enrichment score for each pathway. Dot size reflects adjusted pvalue.

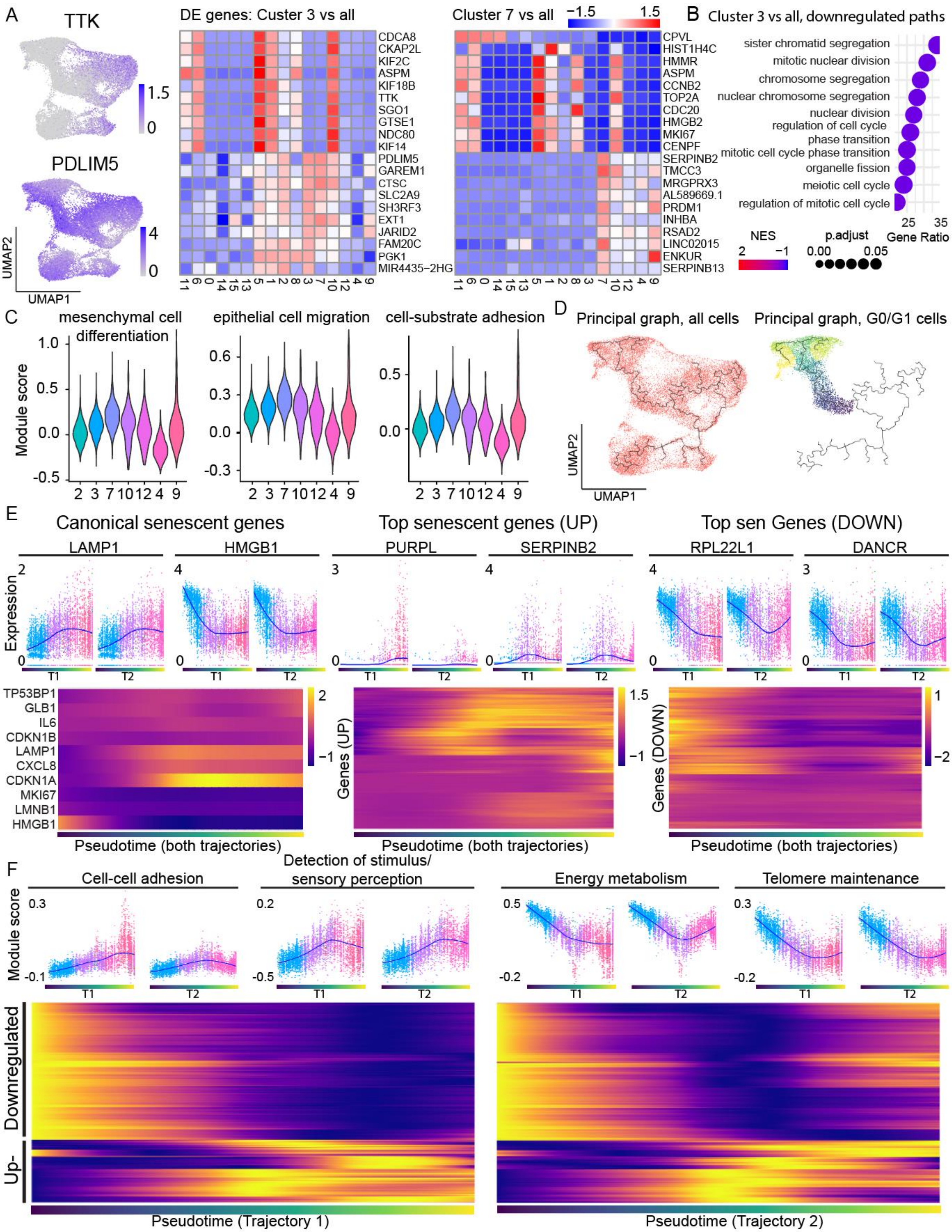
A
TTK
PDLIM5
UMAP2
UMAP1
DE genes: Custer 3 vs all
CDCA8
CKAP2L
KIF2C
ASPM
KIF18B
TTK
SGO1
GTSE1
NDC80
KIF14
PDLIM5
GAREM1
CTSC
SLC2A9
SH3RF3
EXT1
JARID2
FAM20C
PGK1
MIR4435−2HG
11 6 0 14 15 13 5 1 2 8 3 7 10 12 4 9
Cluster 7 vs all
−1.5
1.5
CPVL
HIST1H4C
HMMR
ASPM
CCNB2
TOP2A
CDC20
HMGB2
MKI67
CENPF
SERPINB2
TMCC3
MRGPRX3
AL589669.1
PRDM1
INHBA
RSAD2
LINC02015
ENKUR
SERPINB13
B
Cluster 3 vs all, downregulated paths
sister chromatid segregation
mitotic nuclear division
chromosome segregation
nuclear chromosome segregation
nuclear division
regulation of cell cycle
phase transition
mitotic cell cycle phase transition
organelle fission
meiotic cell cycle
regulation of mitotic cell cycle
25 35
Gene Ratio
NES
2 −1
p.adjust
0.00 0.05
C
mesenchymal cell differentiation
epithelial cell migration
cell-substrate adhesion
Module score
2 3 7 10 12 4 9
D
Principal graph, all cells
Principal graph, G0/G1 cells
UMAP2
UMAP1
E
Canonical senescent genes
Top senescent genes (UP)
Top sen Genes (DOWN)
LAMP1
HMGB1
PURPL
SERPINB2
RPL22L1
DANCR
Expression
T1
T2
TP53BP1
GLB1
IL6
CDKN1B
LAMP1
CXCL8
CDKN1A
MKI67
LMNB1
HMGB1
Genes (UP)
Genes (DOWN)
Pseudotime (both trajectories)
F
Cell-cell adhesion
Detection of stimulus/
sensory perception
Energy metabolism
Telomere maintenance
Module score
Downregulated
Up-
Pseudotime (Trajectory 1)
Pseudotime (Trajectory 2)

**Supplemental Figure 4. Pseudotime trajectory analysis of the quiescence-senescence continuum after etoposide. (A) Left:** Example top up-regulated (bottom) and down-regulated (top) genes for cluster 3 vs. all other clusters (left). UMAPs of top up- and down-regulated genes for cluster 7 can be found in Fig. 2 and S2. Color corresponds to normalized scaled expression. **Right:** Heatmaps of top 10 DE genes (down-regulated at the top and up-regulated at the bottom) for cluster 3 (left) and cluster 7 (right) vs. all cells. Color is average expression in each cluster. **(B)** Gene set enrichment analysis (GSEA) for cluster 3 vs. all other cells. Top 10 pathways (by p.adjusted) upregulated shown. There was only one downregulated pathway, “enamel mineralization”, and it is not shown (Supplementary table 1). Gene ratio plotted on the x-axis. Dots colored by the normalized enrichment score for each pathway. Dot size reflects adjusted pvalue. **(C)** Distributions of module scores for leading-edge genes for top DE pathways between cluster 7 and all other senescent clusters. Violins colored by cluster. **(D)** Pseudotime trajectory analysis using Monocle3. Black line represents the principal graph. All cells were used to generate the trajectory (left) and the branch corresponding to G0/G1 cells (right) was subsetted to exclude cycling cells from downstream analysis. **(E) Top:** Normalized scaled expression of example canonical, top upregulated, or top downregulated senescent genes plotted vs pseudotime. Trajectory 1 and Trajectory 2 analyzed as separate branches. Dot colors correspond to cluster. **Bottom:** Left: Heatmap of 10 canonical senescent genes. Middle: Top 50 up-regulated genes from each senescent cluster over pseudotime. Right: top 50 down-regulated senescent genes from each senescent cluster over pseudotime. Color corresponds to normalized scaled expression. Pseudotime branches combined for simplicity. **(F) Top:** Module score for leading-edge genes of example general biological function groups plotted over pseudotime. Trajectory 1 and Trajectory 2 analyzed as separate branches. Dot colors correspond to cluster. **Bottom:** Heatmap of every differentially expressed GO biological process hit from all senescent clusters over pseudotime. Color represents the Loess fit value for each module score across the pseudotime scatter plots for the pathways listed along the y-axis. Module scores were rescaled for visualization on the same scale.

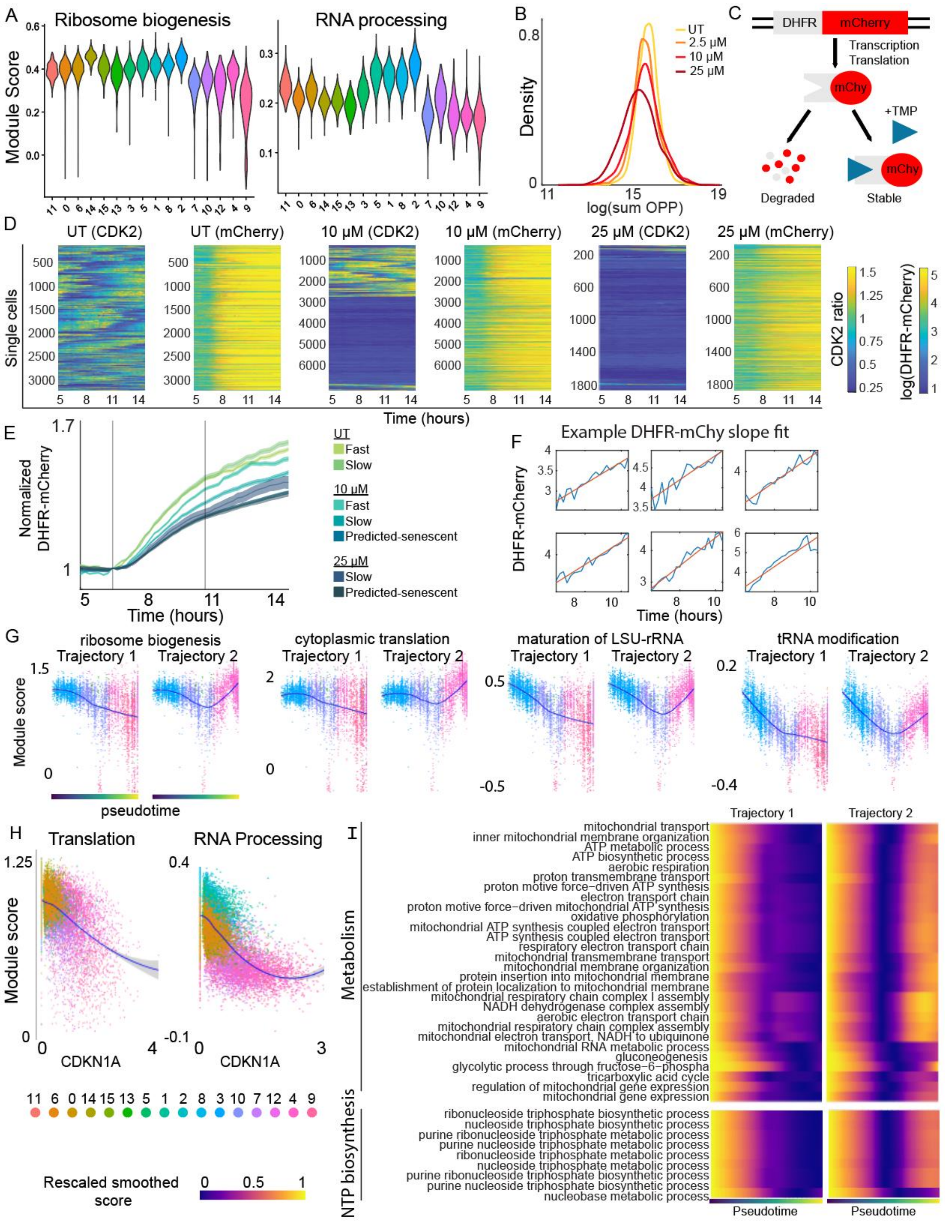

A
Ribosome biogenesis
RNA processing
Module Score
11 0 6 14 15 13 3 5 1 8 2 7 10 12 4 9
B
Density
log(sum OPP)
UT
2.5 μM
10 μM
25 μM
C
DHFR
mCherry
Transcription
Translation
mChy
+TMP
Degraded
Stable
D
UT (CDK2)
UT (mCherry)
10 μM (CDK2)
10 μM (mCherry)
25 μM (CDK2)
25 μM (mCherry)
Single cells
Time (hours)
CDK2 ratio
log(DHFR-mCherry)
E
Normalized DHFR-mCherry
Time (hours)
UT
Fast
Slow
10 μM
Fast
Slow
Predicted-senescent
25 μM
Slow
Predicted-senescent
F
Example DHFR-mChy slope fit
DHFR-mCherry
Time (hours)
G
ribosome biogenesis
cytoplasmic translation
maturation of LSU-rRNA
tRNA modification
Trajectory 1
Trajectory 2
Module score
pseudotime
H
Translation
RNA Processing
Module score
CDKN1A
11 6 0 14 15 13 5 1 2 8 3 10 7 12 4 9
Rescaled smoothed score
I
Trajectory 1
Trajectory 2
Metabolism
mitochondrial transport
inner mitochondrial membrane organization
ATP metabolic process
ATP biosynthetic process
aerobic respiration
proton transmembrane transport
proton motive force−driven ATP synthesis
electron transport chain
proton motive force−driven mitochondrial ATP synthesis
oxidative phosphorylation
mitochondrial ATP synthesis coupled electron transport
ATP synthesis coupled electron transport
respiratory electron transport chain
mitochondrial transmembrane transport
mitochondrial membrane organization
protein insertion into mitochondrial membrane
establishment of protein localization to mitochondrial membrane
mitochondrial respiratory chain complex I assembly
NADH dehydrogenase complex assembly
aerobic electron transport chain
mitochondrial respiratory chain complex assembly
mitochondrial electron transport, NADH to ubiquinone
mitochondrial RNA metabolic process
gluconeogenesis
glycolytic process through fructose−6−phospha
tricarboxylic acid cycle
regulation of mitochondrial gene expression
mitochondrial gene expression
NTP biosynthesis
ribonucleoside triphosphate biosynthetic process
nucleoside triphosphate biosynthetic process
purine ribonucleoside triphosphate metabolic process
purine nucleoside triphosphate metabolic process
ribonucleoside triphosphate metabolic process
nucleoside triphosphate metabolic process
purine ribonucleoside triphosphate biosynthetic process
purine nucleoside triphosphate biosynthetic process
nucleobase metabolic process
Pseudotime

**Supplemental Figure 5. Changes in protein biosynthesis predict cell-cycle fate after etoposide.** **(A)** Distributions of module scores for leading-edge genes belonging to two representative RNA processing and translation-related GO biological processes colored by cluster. **(B)** MCF10A cells were treated with 2.5 μM, 10μM, or 25μM etoposide for 24h followed by a 6d drug-free recovery. Cells were treated with OPP for 30min prior to fixation. Cells were stained for total protein to generate a whole cell mask and OPP was visualized. Sum of the OPP intensity inside the cell mask was plotted as a histogram. Senescent cells increase in area, and it is not clear if the total translation (here) or the translation normalized by area (Fig. 5A) is more biologically important so we quantified both. **(C)** Schematic of the DHFR-mCherry live-cell translation reporter. The DHFR degron is constitutively degraded in the absence of small-molecule TMP but stabilized upon TMP addition. **(D)** Single-cell heatmaps of CDK2 activity and DHFR-mCherry fluorescent intensity (A.U.) over a range of 10hr after TMP addition at 7hr. **(E)** MCF10A cells expressing the CDK2 activity reporter and the DHFR-mCherry translation reporter were imaged by live-cell microscopy for 96h from 6d-10d post etoposide release. The mCherry signal (each cell normalized to its average signal before TMP) was plotted for fast-cycling, slow-cycling, or predicted-senescent cells for each dose. Same data as in the main figure but here, all traces were overlaid for a direct visual comparison. Frames 30-70 were plotted, after which the mCherry signal saturates. TMP was added at the first grey bar. Grey bars bracket the frames used to calculate the mCherry signal slope. **(F)** Example traces with slope calculated by fitting a linear line (orange) through the mCherry intensity (blue) immediately after TMP addition. **(G)** Module scores for leading-edge genes from example RNA processing and translation-related GO biological processes that show different behavior over Trajectory 1 and Trajectory 2 of pseudotime. Dots colored by cluster. Loess fit represented by blue line. **(H)** Module scores for leading-edge genes from RNA processing and translation-related GO biological processes vs. CDKN1A expression. Dots colored by cluster. Loess fit represented by blue line. **(I)** Module scores for metabolism and NTP biosynthesis-related GO biological processes over pseudotime. Color represents the Loess fit value for each module score across the pseudotime scatter plots for the pathways listed along the y-axis. Module scores were rescaled for visualization on the same scale and clustered by similarity.

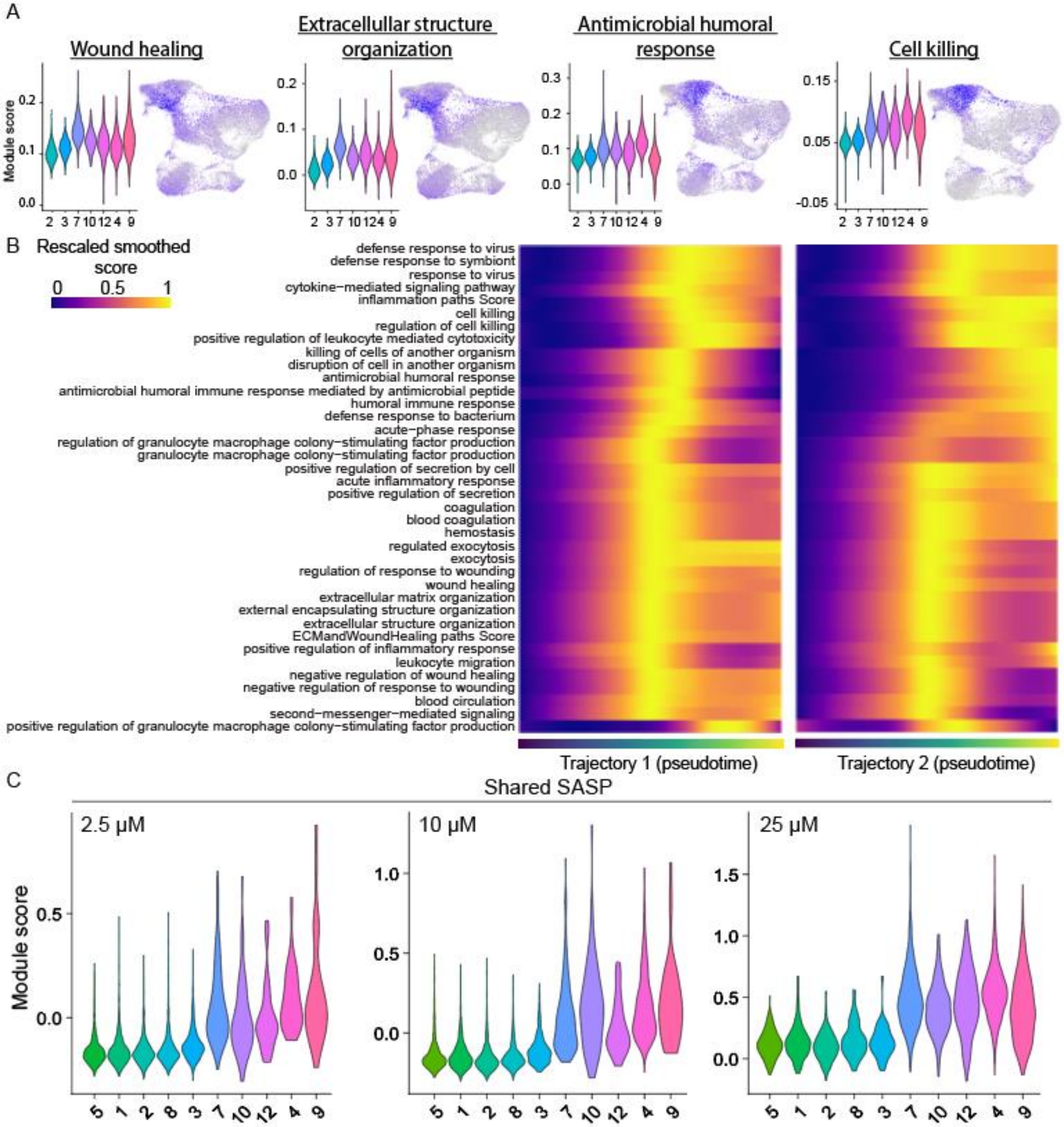


**Supplemental Figure 6. The senescence-associated secretory phenotype is not a general feature of all senescent cells after etoposide. (A)** Distributions of module scores for leading-edge genes belonging to ECM-related or defense response-related GO biological processes colored by cluster and corresponding UMAPs colored by module score. **(B)** Module scores for SASP-related GO biological processes over pseudotime. Color represents the Loess fit value for each module score across the pseudotime scatter plots for the pathways listed along the y-axis. Module scores were rescaled for visualization on the same scale and clustered by similarity. **(C)** Module scores for cells in each etoposide dose group based on expression of shared SASP genes. Shared SASP genes are not uniquely expressed in 25µM cells.

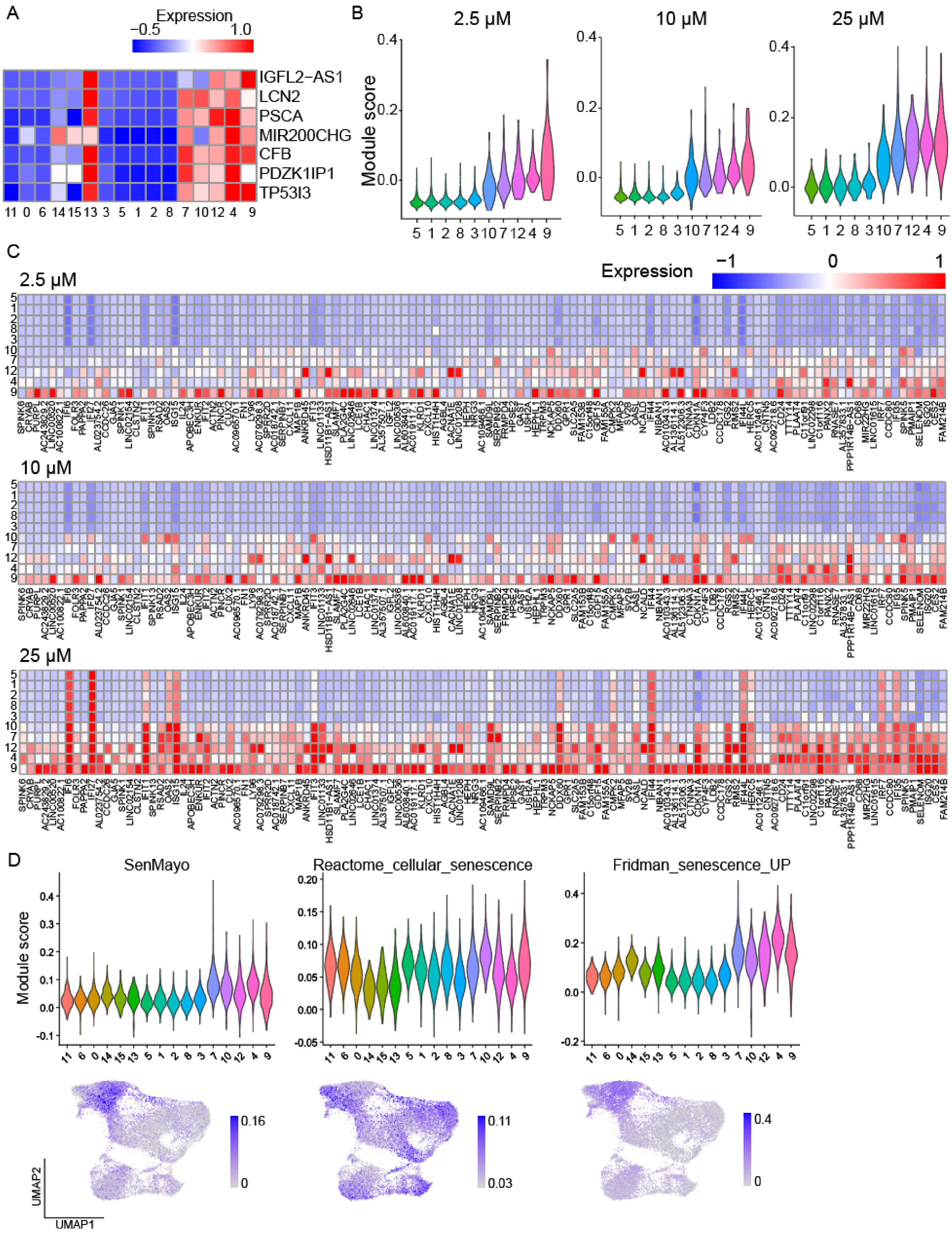
A
Expression
−0.5
1.0
IGFL2−AS1
LCN2
PSCA
MIR200CHG
CFB
PDZK1IP1
TP53I3
11 0 6 14 15 13 3 5 1 2 8 7 10 12 4 9
B
2.5 μM
10 μM
25 μM
Module score
5 1 2 8 3 10 7 12 4 9
C
2.5 μM
10 μM
25 μM
Expression
−1
0
1
D
SenMayo
Reactome_cellular_senescence
Fridman_senescence_UP
Module score
11 6 0 14 15 13 5 1 2 8 3 7 10 12 4 9
0.16
0
0.11
0.03
0.4
0
UMAP2
UMAP1

**Supplemental Figure 7. Genes comprising our senescent gene set are not dose-dependent. (A)** Genes removed from the senescent gene set (see methods) because of significant overlap in expression with untreated spontaneously quiescent cells (cluster 13). **(B)** Module scores for cells in each dose group based on expression of genes in our senescent gene set. Importantly, the gene set is not etoposide dose-dependent and genes are not uniquely expressed in 25μM cells. Only clusters for the treated island of the UMAP are shown. **(C)** Average gene expression (color) per cluster of all genes in the gene list. Cells treated with each dose of etoposide are visualized separately. Only clusters for the treated island of the UMAP are shown. (D) Module scores based on existing senescent gene sets plotted as violins for cells in each cluster as well as single-cell module scores on the UMAPs. **Left:** SenMayo[62], a senescent gene set validated in aged cohort samples across multiple tissues. **Middle:** Reactome_cellular_senescence[45], a curated senescence gene set from the reactome data base accessed through MSigDB[29]. **Right:** Fridman_senescence_UP[63], a curated senescence gene set generated from known senescence markers in the literature accessed through MSigDB[29].
